# Genetic analysis of maize seedling root traits under chilling highlights their importance for early field development

**DOI:** 10.64898/2026.06.11.731510

**Authors:** Fabio Guffanti, Kerstin A. Nagel, Anna Galinski, Carmen Müller, Shree R. Pariyar, Daniela Scheuermann, Claude Urbany, Thomas Presterl, Milena Ouzunova, Chris-Carolin Schön

**Author notes:** Corresponding author: Chris-Carolin Schön.

## Abstract

Characterizing the genetic basis of root system architecture and its role in early plant development is essential for developing maize varieties with improved nutrient uptake, enhanced early vigour, and higher yield potential in temperate regions.

Landraces represent an invaluable source of allelic diversity that can be leveraged to enrich the genetic basis of modern breeding material. In this study, we used a high throughput phenotyping platform to characterize genetic variation for seedling root traits under chilling conditions relevant for early plant establishment in a large doubled haploid (DH) library derived from two European maize landraces.

We dissected the quantitative genetic architecture of twelve seedling root traits using a haplotype-based genome-wide association study, identifying large-effect haplotypes specific to the individual landraces as well as numerous small-effect haplotypes present in both landraces. We validated the effects of four QTL in a biparental population, demonstrating their stability across genetic backgrounds.

We found highly significant correlations between haplotype effects on seedling root traits evaluated in the phenotyping platform and early plant height evaluated in multi environment field trials, demonstrating the relevance of seedling root architecture for early plant establishment. In particular, haplotypes associated with seminal and lateral root length were the major determinants of early plant height under field conditions. Several of the haplotypes increasing seedling root length were absent from a broad panel of flint breeding lines, highlighting their potential as targets for introgression to improve early plant establishment under temperate growing conditions.

**Key message:** Seedling root QTL discovered in a high throughput phenotyping platform under chilling conditions influence early plant development in the field.

## Introduction

With a production exceeding 1.2 billion tons in 2024, maize is one of the most important crops sustaining the global demand for food, animal feed and bioenergy (Food and Agriculture Organization of the United Nations 2026). Its productivity is constrained by multiple abiotic stresses, with the root system playing a pivotal role in mitigating their impacts (Jafari et al. 2024; Lynch 2022). However, despite their recognised importance, roots have been less studied than other plant organs, mainly due to challenges in phenotyping and their strong environmental plasticity (Hochholdinger et al. 2018).

The maize root system consists of embryonic roots (primary and seminal roots), initiated in the embryo, and post-embryonic shoot-borne and lateral roots, initiated after seed germination (Hochholdinger et al. 2004). Embryonic axial roots and early post-embryonic lateral roots are particularly important for early seedling establishment, providing water and nutrients to the developing shoot (Hochholdinger and Tuberosa 2009). In maize, this early developmental stage is highly sensitive to chilling stress, a major constraint in temperate regions, which affects photosynthesis and photoprotection, triggers transcriptomic and hormonal responses, and impairs plant development (Burnett and Kromdijk 2022). Cold tolerant maize, with a root system capable of sustaining early shoot development under chilling conditions, is therefore instrumental for reducing nitrogen losses and improve yields in temperate regions (Ojeda-Rivera et al. 2025).

Under chilling conditions, embryonic roots exhibit a reduced growth rate, decreased total root length, and lower hydraulic conductance, resulting in diminished water and nutrient supply to the shoot (Engels and Marschner 1990; Melkonian et al. 2004; Nagel et al. 2009). Multiple studies indicate that investment in root system expansion is a key feature of cold tolerant, early vigorous maize genotypes (Aroca et al. 2001; Peter et al. 2009; Richner et al. 1996). In seedlings, lateral roots account for a significant proportion of total root length, while representing only a small fraction of root dry weight, making them an efficient means to increase the root surface area (Hund et al. 2004). Colocalizing quantitative trait loci (QTL) and significant phenotypic correlations between root and shoot traits in maize seedlings suggest that longer lateral roots are associated with higher germination speed and confer a functional advantage by supporting higher photosynthetic performance under chilling conditions (Hund et al. 2004; Hund et al. 2007). Seminal roots, a root type that evolved in maize after its divergence from sorghum (Salvi 2017; Singh et al. 2010), have also been shown to enhance early nitrogen and phosphorus uptake (Perkins and Lynch 2021; Zhu et al. 2006). Their positive impact on early nutrient acquisition likely explains why maize domestication favoured an increased seminal root number. However, in drought-prone environments their negative impact on water stress tolerance has favoured a secondary reduction, highlighting the context-dependent relationship between seminal root number and plant productivity (Yu et al. 2024).

In an association study of temperate maize, Reimer et al. (2013) showed that both axial and lateral root elongation respond more strongly to chilling than to heat stress, with maize inbred lines from the flint heterotic pool contributing alleles increasing root biomass accumulation under chilling conditions and dent lines contributing alleles that increase axial root length under optimal temperature conditions. Consistent with these findings, Revilla et al. (2016) highlighted flint lines as a valuable source of alleles for chilling tolerance. As the European flint breeding pool was established by intercrossing a rather small number of founder lines (Messmer et al. 1992), flint landraces may offer opportunities to broaden the genetic basis of modern breeding material. Mayer et al. 2020 could show that doubled haploid lines derived from three European flint landraces carried favourable haplotypes for early plant development that were absent in a broad panel of elite germplasm. In the present study, we (i) characterized the root system architecture of landrace-derived DH lines at the seedling stage under chilling conditions using a high throughput, high precision phenotyping platform to identify heritable root traits, (ii) elucidated their genetic architecture, (iii) linked them to early plant development in the field and (iv) evaluated the potential of introgressing landrace-derived haplotypes into breeding material to improve root system architecture during early stages of maize development.

## Material and Methods

### Background

As part of a comprehensive study on harnessing the genetic diversity of maize landraces, the three landraces Kemater Landmais Gelb (KE, Austria), Petkuser Ferdinand Rot (PE, Germany), and Lalin (LL, Spain), were selected to represent the genetic diversity of 35 European maize landraces (Mayer et al. 2017). A total of 1015 DH lines were derived from these landraces (516 KE, 432 PE, 67 LL) with the in-vivo haploid induction method (Röber et al. 2005) and propagated by self-pollination. 899 DH lines were genotyped with the 600k Affymetrix® Axiom® Maize Genotyping Array, hereafter referred to as the 600k array (Unterseer et al. 2014), and phenotyped for agronomic traits in up to 11 field environments as lines per se and testcrosses (Hoelker et al. 2019). Filtered and imputed genotypic data as well as adjusted means of lines per se within and across environments were obtained from Mayer et al. (2022) for nine agronomic traits including early vigour (at stages V4 and V6), early plant height (at stages V4 and V6), final plant height, anthesis, silking, lodging, and tillering. In addition, adjusted means of a subset of 356 DH lines evaluated for total dry matter testcross yield (TDMY) in two environments were obtained from Hölker et al. (2022).

### Plant material

In this study, 925 DH lines (485 KE, 419 PE, and 21 LL) were phenotyped at the seedling stage up to developmental stage V2 for seedling root traits (experiment 1, E1) with the aim to map quantitative trait loci (QTL) in a haplotype-based genome-wide association study (GWAS). In a second experiment (E2), 94 F2:3 recombinant inbred lines (RILs) derived from a biparental cross were also phenotyped at the same seedling stage with the aim to validate the effects of the haplotypes identified by GWAS. The parental lines of this cross were two DH lines from Kemater, KE0413 (P1) and KE0113 (P2), selected for high background genomic similarity (85% of SNPs from the 600k array were monomorphic) while differing in 26 QTL significantly associated with seedling root traits identified in the GWAS (E1). KE0413 carried 12 QTL alleles with significant trait increasing effects across seven seedling root traits (SRL, LRL, TRL, RSW, CHA, RGR, and RDW, see **Table 1** for trait abbreviations) and one with trait decreasing effect on BAL. In contrast, KE0113 carried eight QTL alleles with significant trait decreasing effects across seven seedling root traits (PRL, SRL, RSD, RSW, CHA, BAL, and SpRL) and five with trait increasing effects across five seedling root traits (PRL, SRL, RSD, CHA, and BAL). The segregating F2 population comprised 554 individuals, from which the 94 F2:3 RILs evaluated in E2 were derived by self-pollination. F2 individuals were genotyped with a custom 15k SNP Illumina array proprietary to KWS SAAT SE & Co. KGaA (hereafter referred to as 15k SNP array).

**Table 1.**
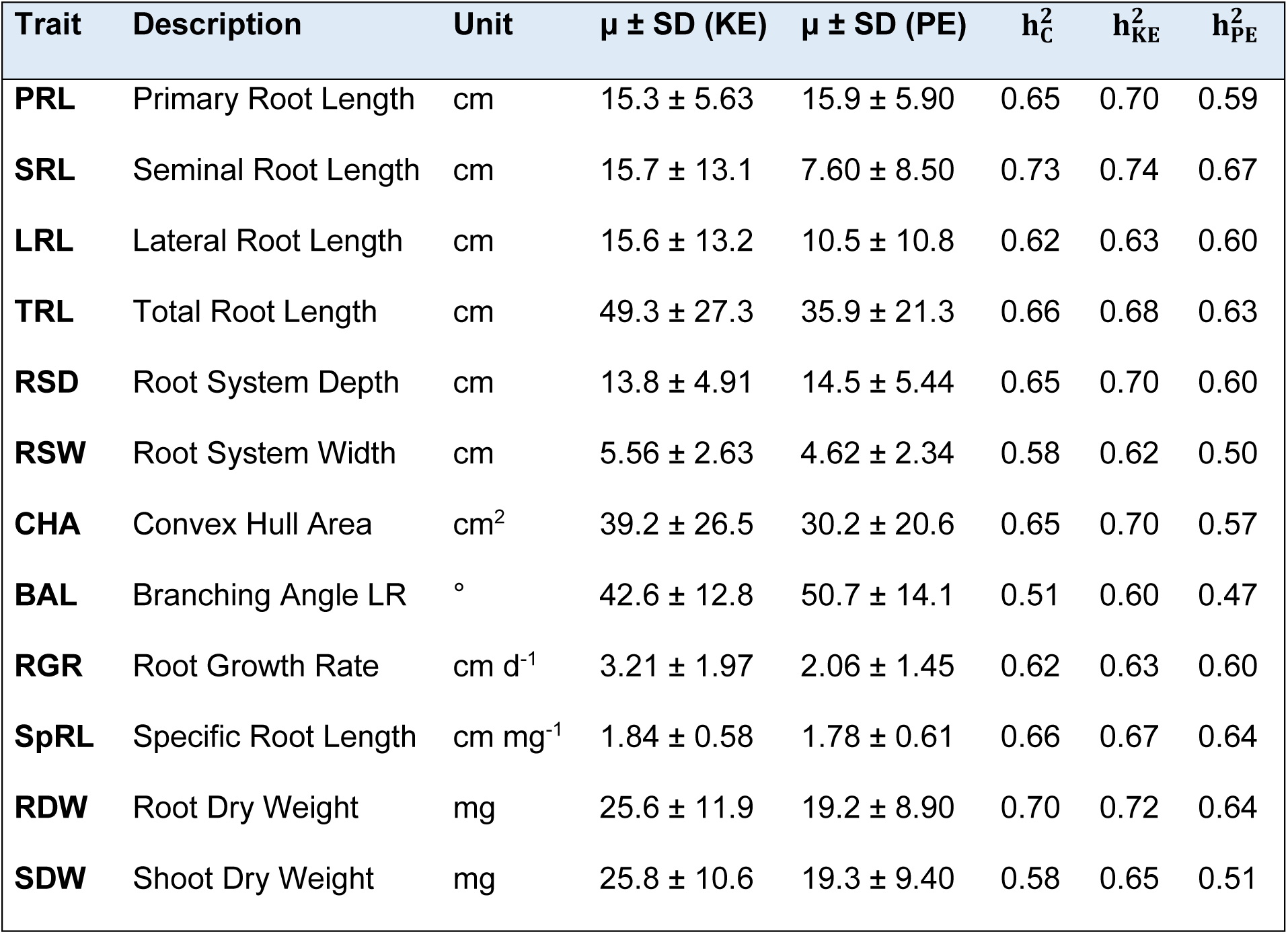
Summary information of the seedling root traits assessed in the phenotyping platform. Indicated are the Trait code, Description, Unit of measure, mean value (μ) and standard deviation (SD) of the adjusted means calculated within doubled haploid (DH) lines derived from Kemater [μ ± SD (KE)] and Petkuser [μ ± SD (PE)], heritability estimation in the combined DH library 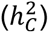, within Kemater 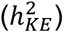 and within Petkuser 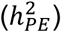

| Trait | Description | Unit | $\mu \pm \text{SD (KE)}$ | $\mu \pm \text{SD (PE)}$ | $h_C^2$ | $h_{KE}^2$ | $h_{PE}^2$ |
| --- | --- | --- | --- | --- | --- | --- | --- |
| **PRL** | Primary Root Length | cm | 15.3 ± 5.63 | 15.9 ± 5.90 | 0.65 | 0.70 | 0.59 |
| **SRL** | Seminal Root Length | cm | 15.7 ± 13.1 | 7.60 ± 8.50 | 0.73 | 0.74 | 0.67 |
| **LRL** | Lateral Root Length | cm | 15.6 ± 13.2 | 10.5 ± 10.8 | 0.62 | 0.63 | 0.60 |
| **TRL** | Total Root Length | cm | 49.3 ± 27.3 | 35.9 ± 21.3 | 0.66 | 0.68 | 0.63 |
| **RSD** | Root System Depth | cm | 13.8 ± 4.91 | 14.5 ± 5.44 | 0.65 | 0.70 | 0.60 |
| **RSW** | Root System Width | cm | 5.56 ± 2.63 | 4.62 ± 2.34 | 0.58 | 0.62 | 0.50 |
| **CHA** | Convex Hull Area | cm<sup>2</sup> | 39.2 ± 26.5 | 30.2 ± 20.6 | 0.65 | 0.70 | 0.57 |
| **BAL** | Branching Angle LR | ° | 42.6 ± 12.8 | 50.7 ± 14.1 | 0.51 | 0.60 | 0.47 |
| **RGR** | Root Growth Rate | cm d<sup>-1</sup> | 3.21 ± 1.97 | 2.06 ± 1.45 | 0.62 | 0.63 | 0.60 |
| **SpRL** | Specific Root Length | cm mg<sup>-1</sup> | 1.84 ± 0.58 | 1.78 ± 0.61 | 0.66 | 0.67 | 0.64 |
| **RDW** | Root Dry Weight | mg | 25.6 ± 11.9 | 19.2 ± 8.90 | 0.70 | 0.72 | 0.64 |
| **SDW** | Shoot Dry Weight | mg | 25.8 ± 10.6 | 19.3 ± 9.40 | 0.58 | 0.65 | 0.51 |

### Phenotypic data collection

In E1, eight replications of each of the 925 DH lines were evaluated using the germination paper-based phenotyping platform GrowScreen-PaGe (Gioia et al. 2017). The experiment was a randomized complete block design (RCBD) with eight blocks. The DH lines were divided into three batches and evaluated across 12 sequential runs in a growth chamber, each run including two replicates of a single batch (600–650 plants per run). In E2, seven replications of 94 F2:3 RILs and 21 replications of P1 and P2 were evaluated in the GrowScreen-PaGe, all in the same growth chamber. The experiment was a RCBD with seven blocks, and the parents were included as triplicate entries.

For experiments E1 and E2, 10 seeds from each genotype were pre-soaked in distilled water for 8 hours at room temperature to promote uniform germination. The seeds were then placed between two round filter papers in a 94 mm Petri dish, to which 3 ml of deionized water was added before sealing with Parafilm. Petri dishes were kept in the dark by putting them into a box and arranged in a growth chamber at 24/22 °C, with a 14/10 h cycle and 65% relative humidity. After approximately 40 hours, seedlings with a visible primary root were transferred onto germination paper, with one seedling per paper, and sprayed with nutrient solution. 50 germination papers were arranged in one growth container, and containers were transferred to the growth chamber. This time point is referred to as 0 days after transplantation (dat). In the growth chamber, water and nutrients were supplied in equal amounts at the bottom of each growth container and absorbed by capillary action by the germination papers. The nutrient solution consisted of a modified Hoagland solution to one-third strength (stock solution, 5mM KNO3, 5mM Ca(NO3)2, 2mM MgSO4, 1mM KH2PO4, plus Fe-EDTA and trace elements) that was replaced after one week (Hoagland and Arnon 1938). Environmental conditions in the growth chamber were maintained at 65% relative air humidity, the day/night cycle was set to 14/10 hours, with a light intensity of 400 μmol m^−2^ s^−1^ supplied during the day with artificial lights. To allow seedling establishment after transplantation onto germination paper, the temperature was maintained at 24/22°C day/night until 7 dat in E1 and until 8 dat in E2. Following this period, temperature was shifted to chilling conditions of 16/12°C day/night for additional 7 days to simulate soil temperature during early growth of maize in temperate regions. Plants were grown until 14 dat (E1) and 15 dat (E2), when they were harvested to measure shoot and root biomass. The root systems were non-destructively imaged at 5, 7, 9, 12, and 14 dat in E1 (**Fig. 1a**) and at 5, 8, and 15 dat in E2. Root traits were assessed as described by Pariyar et al. (2021) using a custom software and manual curation (**Fig. S1**).

**Fig. 1.**
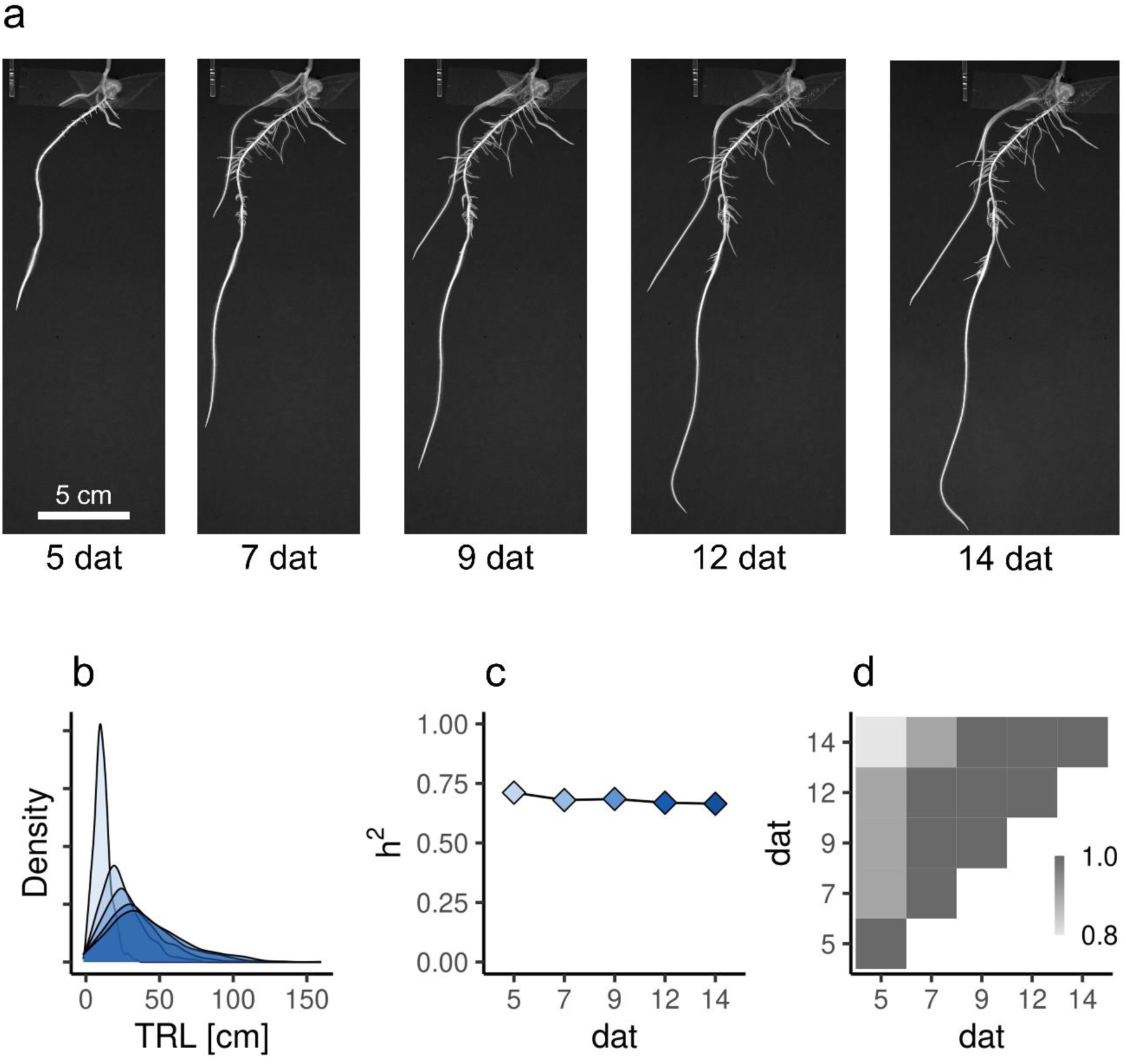
Evaluation of the doubled haploid (DH) library in the GrowScreen-PaGe. a) Representative pictures of a plant at 5, 7, 9, 12, 14 days after transplantation (dat). Seedling root traits were measured at each time point. b) Distributions of adjusted means of total root length (TRL, cm) across five time points, with progressively darker blue tones indicating later time points. c) Heritability estimation of TRL across five time points. d) Pearson correlation coefficients between TRL across five time points, indicated by a grey colour scale

In E1, the measured root traits at each time point included primary root length (PRL, cm), seminal root length (SRL, cm), lateral root length (LRL, cm), crown root length (CRL, cm), total root length (TRL, cm), root system width (RSW, cm), root system depth (RSD, cm), area of the convex hull that encompasses the root system (CHA, cm^2^), and branching angle of lateral roots (BAL, °). At harvest (14 dat), roots and shoots were separated at the seed. Subsequently, root dry weight (RDW, mg) and shoot dry weight (SDW, mg) were determined after oven drying at 65 °C for one week. TRL increased approximately linearly over time for all genotypes (**Fig. S2**). Therefore, the root growth rate between two timepoints (dat) t1 and t2 [RGR_(t,t)_, cm d^−1^], reflecting the speed of root system expansion, was calculated as: 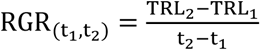. The specific root length (SpRL, cm mg^−1^), reflecting the efficiency of soil exploration, was defined as the ratio between TRL and RDW. In E2, the traits TRL, RSW, RSD, CHA, RDW, SDW, RGR, and SpRL were measured.

In both experiments, a categorical tag indicating normal or abnormal plant growth was assigned to each image. Plants were classified as abnormal and discarded if they exhibited a dead primary root, detachment from the germination paper, improper kernel germination, fungal contamination, or null growth rate between 5 and 7 dat. After filtering, 815 DH lines derived from KE (n = 440) and PE (n = 375) were retained in E1. DH lines from LL were underrepresented and therefore not considered in further analyses. In E2, all genotypes were retained.

### Phenotypic data analysis

All statistical analyses were conducted using R (R Core Team 2025). Data visualization was based on the R package *ggplot2* (Wickham 2016). The statistical model for estimating genotype and error variance components for each seedling root trait in E1 and E2 was:

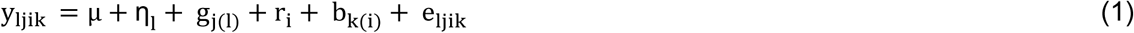

Where y_ljik_ denotes the phenotypic observation of genotype j in group l, experimental run i, and block k for a given trait, μ is the intercept (overall mean); η_l_ is the effect of the group l (with l = 1, 2 for the landraces KE, PE, respectively and l = 3 for the two parental lines in E2); g_j(l)_ is the effect of genotype j nested in group l; r_i_ is the effect of the experimental run i in E1; b_k(i)_ is the effect of block k nested in experimental run i; and e_ljik_ is the residual error.

To estimate variance components, all effects were treated as random and assumed to be independent and identically normally distributed, except g_j(l)_ for l = 3 and η_l_ that were treated as fixed effects. Outliers were defined based on the distribution of model residuals. Observations more than three interquartile ranges below or above the central 50% of the data were classified as outliers and excluded from further analyses. Variance components and their standard errors were estimated using the restricted maximum-likelihood method implemented in ASReml-R package (Butler 2017). Trait heritabilities (h^2^) were calculated on an entry-mean basis according to Holland et al. (2003). Adjusted means were obtained from equation (1), treating g_j(l)_ as fixed effect and excluding the term η_l_. Phenotypic correlations among traits were calculated as the Pearson correlation coefficient (rp) of the adjusted means for pairwise trait combinations. Correlations were considered significant after correcting for multiple testing using a false discovery rate (FDR) of 5% (Benjamini and Hochberg 1995).

In experiment E1, variance components, heritabilities, and phenotypic correlations were also estimated within landraces PE and KE, using only the landrace-specific phenotypic data and dropping the term η_l_ in equation (1).

### Principal Component Analysis

Centered and scaled adjusted means of twelve seedling root traits at 14 dat (E1) (**Table 1**) were used for principal component analysis (PCA). Missing phenotypic values were imputed using the arithmetic mean of the respective trait. PCA was performed using the *prcomp* function in base R, and results were visualized using the R package *factoextra* (Mundt 2026). Differences between KE- and PE-derived DH lines in the PC1-PC2 space were tested using permutational multivariate analysis of variance (PERMANOVA) based on Euclidean distances with 1000 permutations using the *adonis2* function of the R-package *vegan* (Oksanen J et al. 2025).

### Genome wide association study

For GWAS, genotypic data consisted of 496,426 filtered and imputed SNPs mapping unambiguously to the same chromosomes in both B73v4 (Jiao et al. 2017) and B73v5 (Hufford et al. 2021) reference genomes. Following Mayer et al. (2020), we constructed haplotypes by concatenating non-overlapping windows of 10 consecutive SNPs and coded them as bi-allelic markers to be used in the GWAS. If haplotypes were in perfect linkage disequilibrium (LD, r^2^ = 1) one of them was randomly chosen resulting in 102,947 haplotypes with a minor allele count ≥ 3 in the set of 791 DH lines genotyped and phenotyped for root traits (438 derived from KE and 358 from PE). For comparison, we also conducted a GWAS using 125,226 SNPs (r^2^< 1) with a minor allele count ≥ 3 in the same set of 791 DH lines. Phenotypic data consisted of the adjusted means of the twelve seedling root traits at 14 dat (E1) described in **Table 1**.

For each of the twelve seedling root traits, a univariate linear mixed model implemented in GEMMA (Zhou and Stephens 2012) was fitted using the leave-one-chromosome-out (LOCO) approach as described by Yang et al. (2014):

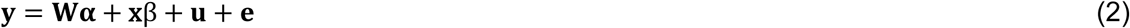

where **y** is the *n-*dimensional vector of the adjusted means for a given trait, with *n* being the number of DH lines, **W** is a (*n x 2*) design matrix assigning the phenotypic values to **α**; **α** is the *two*-dimensional vector of fixed effects (intercept and landrace effect of KE), β is the fixed effect of the tested haplotype, **x** is a *n*-dimensional vector of corresponding genotype scores coded as 0 or 2, **u** is the *n*-dimensional vector of random polygenic effects with **u**∼N(0, **K**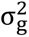); and **e** is the *n*-dimensional vector of random residual errors, with **e**∼N(0, **I**_**n**_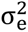). **K** is the (*n x n*) genomic relationship matrix calculated based on the filtered SNP data according to Astle and Balding (2009), excluding the SNPs on the same chromosome as the tested marker. **I**_**n**_ is the (*n x n*) identity matrix. 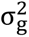 and 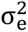 refer to the genetic and residual variance component, respectively. Significance of the haplotype-trait associations was assessed with a Wald test, using a FDR of 5% (Benjamini and Hochberg 1995).

The distribution of association signals was assessed using Manhattan and quantile-quantile (Q-Q) plots (**Fig. S3**). Significant haplotypes in high LD (r^2^ > 0.6) were grouped into genomic regions delimited by the first and last haplotype of the group and represented by the most significant haplotype. For overlapping genomic regions, only the most significant haplotype was retained when fitting a multi-locus QTL model (**Fig. S3**).

### Multi-locus model selection

To build a multi-locus QTL model for each seedling root trait, we conducted a backward elimination of haplotypes associated with the trait under study:

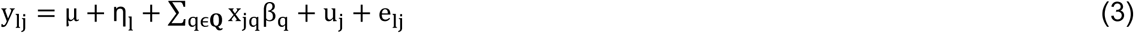

Where y_lj_ is the scaled and centered adjusted mean of line j belonging to landrace l for a given trait; μ is the common intercept; η_l_ is the fixed effect of the landrace l; x_jq_ is the genotype score of line j for haplotype q coded as 0 or 2; β_q_ is the fixed effect of haplotype q, u_j_ is the random polygenic effect of line j, with **u**∼N(0, **K**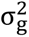); and e_lj_ is the random residual error, with **e**∼N(0, **I**_**n**_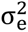). **K** is the (*n x n*) genomic relationship matrix calculated from filtered SNPs according to Astle and Balding (2009). Significance of fixed effects was tested with a Wald test implemented in ASReml-R. At each step of backward elimination, the significance of a given haplotype was tested when entering the model last. The least significant haplotype was removed from the model if P > 0.01, until all haplotypes included in the model were significant. After stepwise backward elimination, **Q** denotes the final set of retained haplotypes representing genomic regions with significant effects on the target trait (hereafter referred to as lead haplotypes).

Haplotypes located within 1 Mb and in high LD (r² > 0.6) with the lead haplotypes, regardless of their statistical significance, were considered part of the associated genomic regions (QTL), with the first and last such haplotypes defining the boundaries of the QTL confidence interval. QTL associated with different traits were considered overlapping when their confidence intervals overlapped and the lead haplotypes were in high LD (r² > 0.6). QTL were named following McCouch (1997), using the prefix “q” followed by the trait code, chromosome number, and a sequential capital letter indicating their order along the chromosome.

### Estimation of variance explained by lead haplotypes

To estimate the proportion of genetic variance explained by the lead haplotypes associated with a given seedling root trait we used the following statistical model:

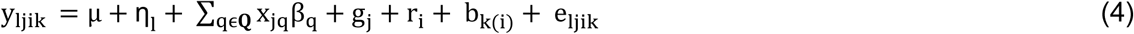

Where y_ljik_ denotes the phenotypic observation of genotype j in group l, experimental run i, and block k for a given trait; μ is the intercept (overall mean); η_l_ is the fixed effect of the group l (with l = 1, 2 for the landraces KE, PE, respectively); x_jq_ is the genotype score of line j for haplotype q coded as 0 or 2; β_q_ is the fixed effect of haplotype q, with **Q** representing the set of lead haplotypes; g_j_ is the random effect of genotype j; r_i_ is the random effect of the experimental run i; b_k(i)_ is the random effect of block k nested in experimental run i; and e_ljik_ is the random residual error. Random effects were assumed to be independent and identically normally distributed. The proportion of genetic variance explained simultaneously by all lead haplotypes in **Q** was estimated as the reduction in genetic variance when haplotype effects were included compared to the model with **Q** = ∅.

For agronomic traits, this proportion was estimated using environment-specific adjusted means as the response variable in a model including landrace, haplotype, environment, and haplotype-by-environment interaction as fixed effects, and genotype as a random effect, following Mayer et al. (2020).

### Validation of lead haplotypes in the biparental population

To estimate the effects of the lead haplotypes associated with a given seedling root trait in the biparental population, we performed seedling root trait-specific multiple marker regression in the F2:3 RILs, with each lead haplotype represented by the closest SNP genotyped in the biparental population. We used genotypic data from the 15K array for the 554 F2 lines after excluding SNPs not represented on the 600K array, those with missing calls in P1 or P2, heterozygosity in P1 or P2, >5% missing data, or >5% non-parental alleles. Phenotypic data consisted of adjusted means of F2:3 RILs. The statistical model was as described for model (3) but excluding the term u_j_. In this model, y_lj_ denotes the adjusted mean of line j belonging to group l (l = 1 for parents, l = 2 for F2:3 RILs) for a given trait; η_l_ represents the effect of group l; x_jq_ is the genotype score of line j for the SNP q closest to the respective lead haplotype, coded as 0, 1, 2 for the P1, heterozygous, and P2 genotype, respectively; β_q_ is the fixed effect of SNP q, with **Q** representing the set of SNPs representing the respective lead haplotypes; and e_lj_ is the random residual error. Significance of fixed effects was tested with a Wald test implemented in ASReml-R.

### Estimation of haplotype effects on root and agronomic traits

To assess the simultaneous effect of each lead haplotype on seedling root and agronomic traits, we extended model (2) to a bivariate model for all seedling root-agronomic trait pairs:

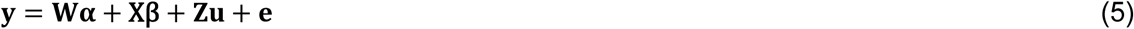

where **y** is the *2n-*dimensional vector of scaled and centered adjusted means for a given seedling root and agronomic trait combination, with *n* denoting the number of lines phenotyped for both traits; **W** is a (*2n x 4*) design matrix assigning phenotypic values to fixed effects; **α** is a *four*-dimensional vector of trait-specific fixed intercepts and landrace effects; **X** is a (*2n x 2*) matrix of genotype scores coded as 0 and 2; **β** is a *two*-dimensional vector of trait-specific fixed haplotype effects; **Z** is an incidence matrix (with **Z** = **I**_**2n**_) assigning observations to genotypes; **u** is the *2n*-dimensional vector of trait-specific random polygenic effects, with **u**∼**N**(**0**, **G** ⊗ **K**); and **e** is the *2n*-dimensional vector of random residual errors, with **e**∼**N**(**0, E** ⊗ **I**_**n**_). **K** is the (*n x n*) genomic relationship matrix calculated from filtered SNPs according to Astle and Balding (2009). **I**_n_ is the (*n x n*) identity matrix. 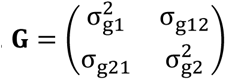 and 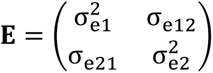 are the genetic and error variance-covariance matrices for a seedling root and agronomic trait combination, and ⊗ denotes the Kronecker product. Lead haplotypes were considered significantly associated with both traits when the two-sided 95% confidence interval of their effect estimates did not include zero for both traits.

### Assessing the frequency of lead haplotypes in breeding lines

To assess the frequency of the identified lead haplotypes in breeding lines, we obtained filtered and imputed genotypic data of 65 flint inbred lines described in Mayer et al. (2020). Analogously to the procedure used for the DH library, we constructed haplotypes based on 496,426 SNPs mapping unambiguously to B73v4 and B73v5, retained the 102,947 haplotypes used in the GWAS, and calculated their frequencies in the panel of 65 breeding lines.

## Results

### Heritability of seedling root traits in landrace-derived DH lines

We evaluated seedling root system development in 815 landrace-derived DH lines under chilling conditions by quantifying the length of embryonic and early post-embryonic roots across five time points until developmental stage V2 (14 dat) (**Fig. 1a**). Phenotypic variance of total root length (TRL) progressively increased over time (**Fig. 1b**), while the trait heritability remained stable (>0.66) due to the parallel increase in genetic and residual variance (**Fig. 1c**). TRL measurements were strongly correlated across time points (rp > 0.8, **Fig. 1d**).

A similar pattern was observed for the other seven seedling root traits measured repeatedly over time. Therefore, traits measured at the final time point (14 dat) were considered representative of the seedling root system architecture, as seminal and lateral roots were more developed at this stage and biomass-related traits were also assessed at the same time point. In the following, we focus on ten root traits with heritabilities ranging from 0.51 to 0.73 and on the root growth rate in chilling conditions between 7 and 14 dat (RGR, h^2^ = 0.62) to characterize the seedling root system architecture (**Table 1**). In addition, shoot dry weight (SDW; h² = 0.58) was included in the analysis to evaluate its relationship with root architectural traits. Crown root length and the length of lateral roots originating from seminal roots were not considered further, as 75% and 91% of the measured plants did not develop them.

### Correlating root and agronomic traits

The principal component analysis (PCA) based on the seedling root traits and SDW separated DH lines derived from the two landraces, with the first two principal components explaining together 70% of the total phenotypic variance (**Fig. 2a**). KE-derived DH lines displayed, on average, more vigorous root and shoot systems, characterized by greater TRL (**Fig. 2b**), longer seminal and lateral roots, faster root growth rates, and higher root and shoot dry weight. In contrast, PE-derived DH lines exhibited, on average, slightly deeper root systems (**Table 1**).

**Fig. 2.**
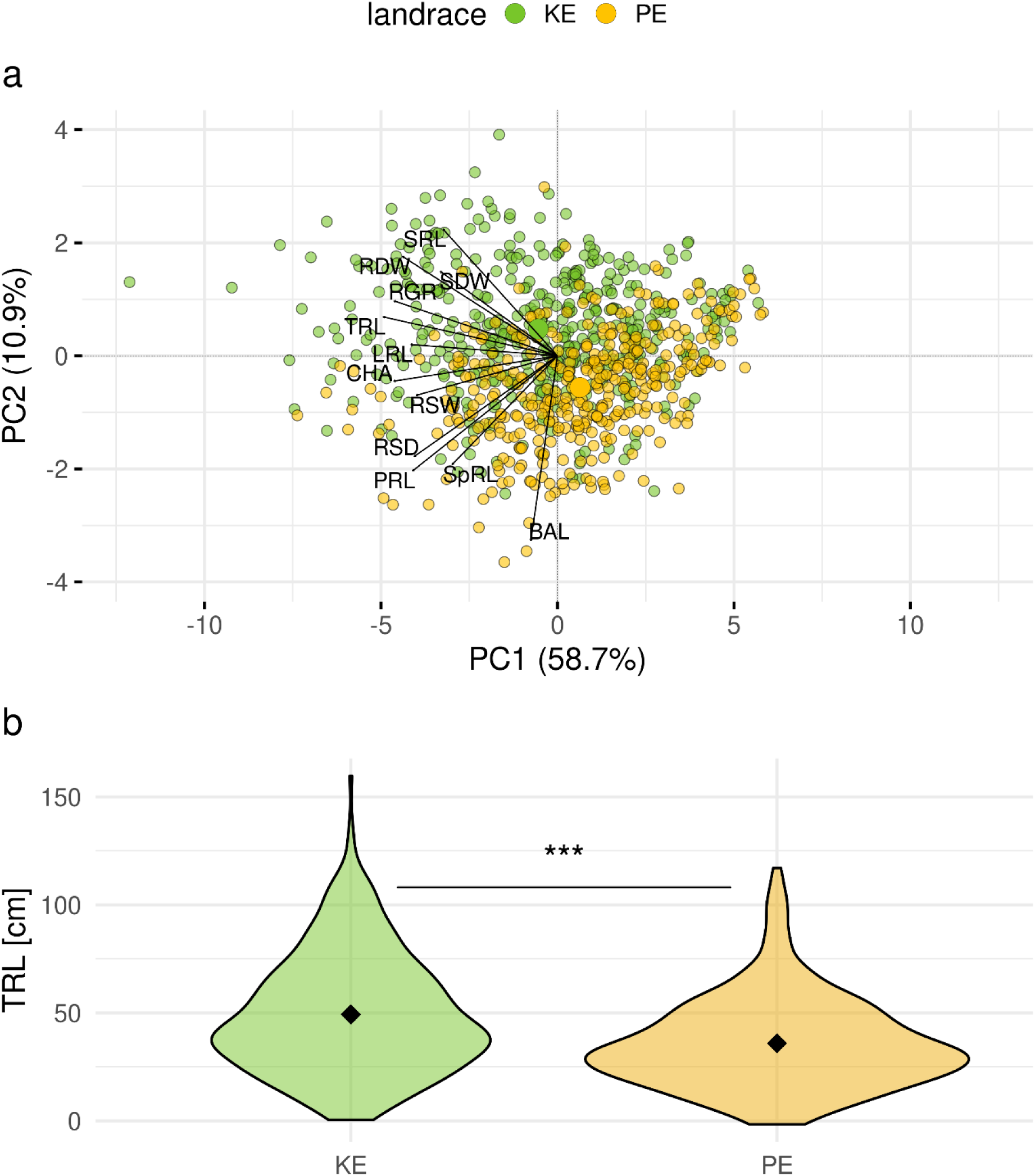
Principal component analysis (PCA) of seedling root traits (SRT). a) PCA biplots based on twelve SRT assessed on 815 doubled haploid (DH) lines. DH lines derived from Kemater (n = 440, KE) and Petkuser (n = 375, PE) are represented as green and orange dots, respectively, with larger dots indicating group centroids. Traits are represented as lines. *x* and *y* axis refer to the first (PC1) and second (PC2) principal component, respectively, with the proportion of total phenotypic variance explained indicated. b) Distributions of adjusted means for total root length (TRL) showing significant differences between KE and PE, as determined by a t-test (*** P < 0.001). Mean values for KE and PE are indicated by diamonds. For trait descriptions, refer to **Table 1**

Phenotypic correlations among traits assessed in the phenotyping platform and in field trials were consistent across PE and KE DH libraries (**Fig. S4**). In both landraces, seedling root traits were positively correlated with each other in almost all cases, with rp ranging from 0.12 to 0.98. Primary root length (PRL), lateral root length (LRL), and seminal root length (SRL) were weakly to moderately correlated (rp < 0.6). Root system depth (RSD) was almost perfectly correlated with PRL (rp ∼ 1) and weakly to moderately with SRL and LRL. Root system width (RSW), convex hull area (CHA), TRL, root dry weight (RDW), and RGR were moderately to strongly correlated with the length of primary, seminal and lateral roots (rp = 0.5-0.8). Specific root length (SpRL) showed weak to moderate correlations with other traits (rp = 0.2-0.6), and the branching angle of lateral roots (BAL) was largely independent of other traits (rp < 0.4). SDW was positively correlated with most root traits (rp = 0.2-0.7).

Most seedling root traits were significantly correlated with early developmental traits in both DH libraries (**Fig. S4**). Exceptions were SRL, which showed significant associations only in KE, and BAL and SpRL, which did not show a significant correlation. Correlations of seedling root traits with other agronomic traits were generally weak, inconsistent, and mostly landrace specific (**Fig. S4**).

### Uncovering the genetic basis of root architectural variation

Based on the GWAS performed in the population of landrace-derived DH lines, we identified 137 significant haplotype-trait associations (HTAs) corresponding to 109 genomic regions carrying QTL for one or several of the twelve seedling root traits (**Fig. 3**). 86 QTL were associated with a single trait, 19 QTL shared between two traits, three QTL shared between three traits, and one QTL was associated with four traits. For these colocalizing QTL, the lead haplotypes were in high LD (r^2^ > 0.6) and showed effect directions consistent with the phenotypic correlations found among traits (**Fig. S5**). From now on we will refer to the 109 genomic regions as QTL irrespective if they carry one or several HTAs.

**Fig. 3.**
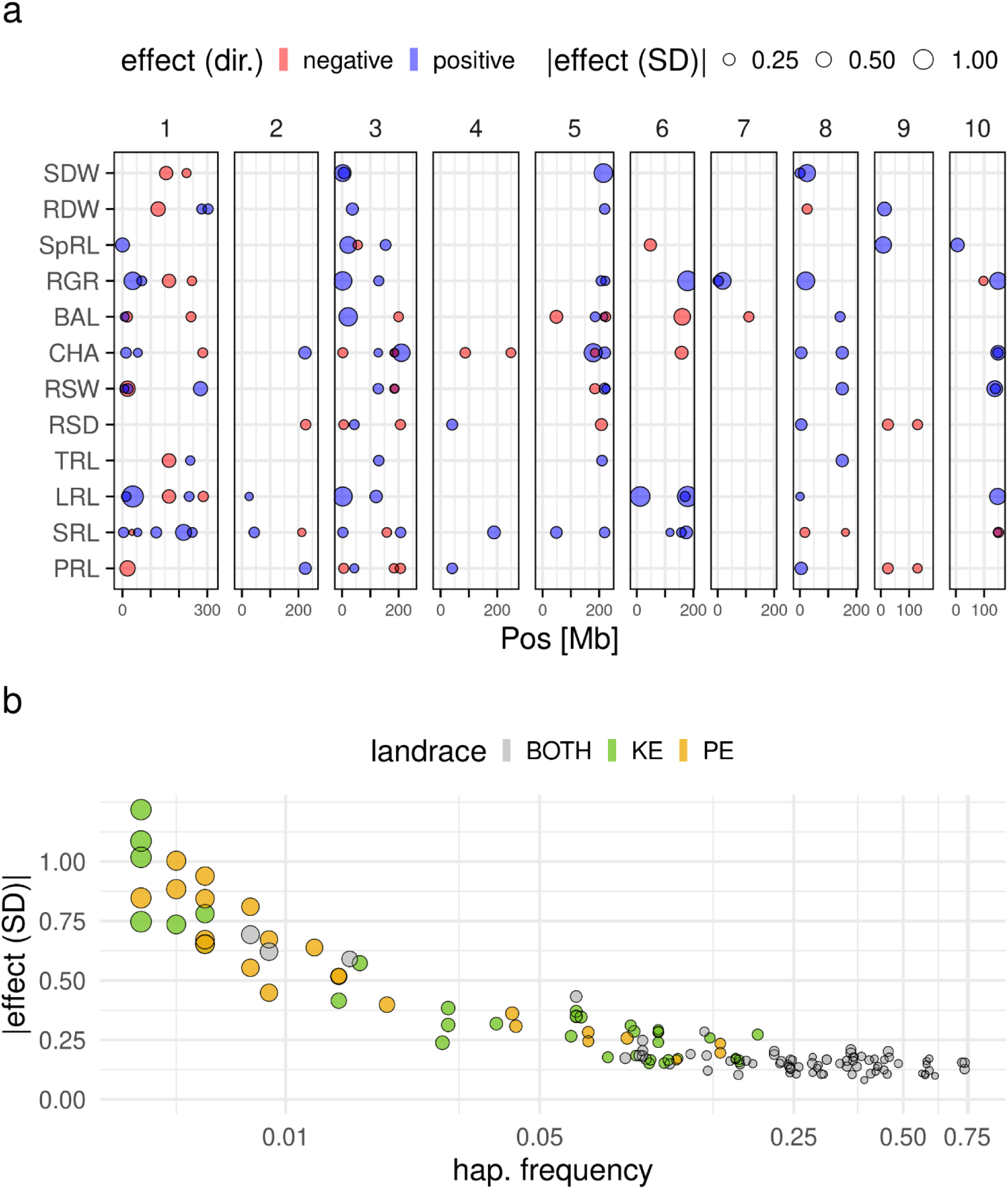
Haplotypes significantly associated with seedling root traits (SRT) identified in the doubled haploid (DH) library. a) Genomic positions of 137 haplotypes identified for twelve SRT (y-axis). Haplotypes are shown as dots across the 10 chromosomes according to their physical positions (Mb) on the x-axis based on B73v5, colored in blue and red if they had positive (trait-increasing) or negative (trait-decreasing) effects, respectively. Dot size is proportional to the absolute additive effect size, expressed in standard deviations of the respective trait [|effect (SD)|]. b) Relationship between haplotype frequency (x-axis) and |effect (SD)| (y-axis) for the 137 haplotypes shown in (a). Dots are colored grey, green, or yellow to indicate haplotypes present in both landraces, specific to Kemater (KE) or Petkuser (PE), respectively, with size proportional to the standard error of the effect estimate. For trait descriptions, refer to **Table 1**

Absolute haplotype effect size decreased with increasing haplotype frequency (**Fig. 3b**). Haplotypes with concordant effects across landraces generally exhibited small effects, whereas haplotypes with moderate to large effects were rare (frequency < 0.05) and predominantly specific to a single landrace (**Fig. 3b**). Considering lead haplotypes present in at least three DH lines per landrace, effect estimates were significantly correlated between KE and PE (n = 66, r = 0.74, P < 0.001) (**Fig. S6**).

Overall, the genetic architecture of the investigated seedling root traits was characterized by a mixture of haplotypes with large (>0.7 SD, 9%), moderate (0.3-0.7 SD, 19%), and small (<0.3 SD, 72%) effect estimates. Depending on the trait, between 19% and 53% of the total genetic variance could be explained by the multi-locus model including all significant lead haplotypes (eq. 4, **Table 2**).

**Table 2.** Summary information of the lead haplotypes associated with seedling root traits identified in the combined doubled haploid library. Indicated are the Trait code, number of lead haplotypes significantly associated with a given trait (No. lead hap.), the minimum and maximum absolute additive haplotype effect expressed in standard deviations of the respective trait [|effect| min (SD), |effect| max (SD)], and the proportion of genetic variance explained by the multi-locus model including all lead haplotypes [GVE (%)]

| Trait | No. lead hap. | \|effect\| min (SD) | \|effect\| max (SD) | GVE (%) |
| --- | --- | --- | --- | --- |
| **PRL** | 10 | 0.13 | 0.52 | 24 |
| **SRL** | 21 | 0.08 | 0.57 | 51 |
| **LRL** | 13 | 0.11 | 1.22 | 39 |
| **TRL** | 5 | 0.14 | 0.37 | 25 |
| **RSD** | 9 | 0.15 | 0.26 | 25 |
| **RSW** | 13 | 0.11 | 0.55 | 45 |
| **CHA** | 19 | 0.10 | 0.81 | 53 |
| **BAL** | 12 | 0.10 | 0.85 | 46 |
| **RGR** | 14 | 0.13 | 1.02 | 39 |
| **SpRL** | 7 | 0.14 | 0.65 | 27 |
| **RDW** | 7 | 0.16 | 0.43 | 23 |
| **SDW** | 7 | 0.13 | 0.88 | 19 |

Of the 109 QTL discovered in the GWAS, 15 QTL were segregating for the traits evaluated in the biparental population. Of the 15 QTL, 11 showed concordant effect directions between the GWAS and biparental population, and four of these QTL were significant in the biparental population (**Fig. 4a**). These comprised QTL on Chr. 1 at 19.7 Mb for RSW (qRSW01C); Chr. 3 at 5.6 Mb for PRL and RSD (qPRL03A, qRSD03A) and at 55.4 Mb for SpRL (qSpRL03B); and Chr. 10 at 148.6 Mb for CHA (qCHA10A) (**Fig. 4b**).

**Fig. 4.**
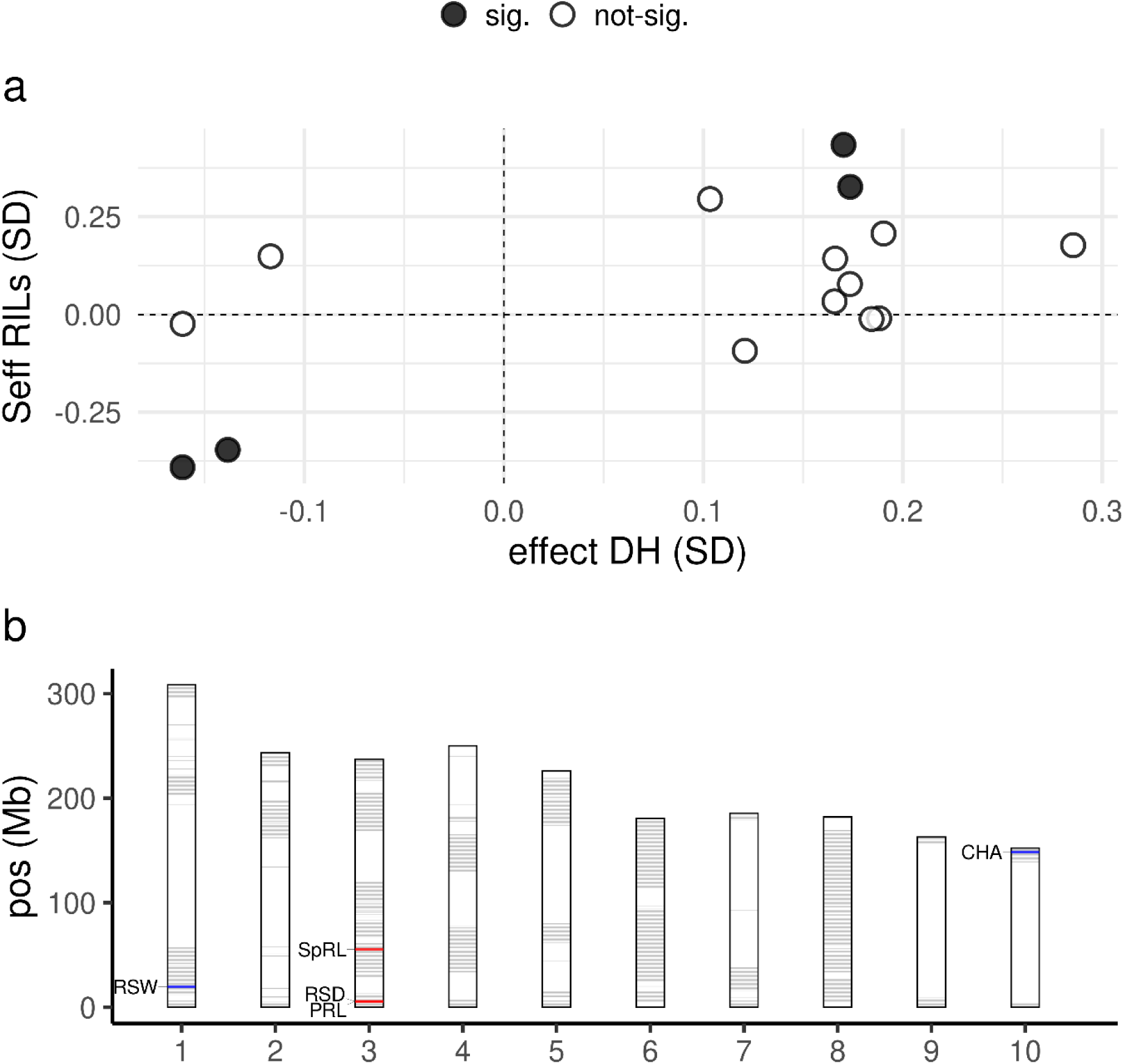
Validation of GWAS-identified haplotypes in a biparental population. a) Scatterplot of the additive effects of 15 lead haplotypes representing QTL identified by GWAS in the DH library (x-axis) and of the corresponding SNPs in the F2:3 RILs (y-axis), expressed in standard deviations of the respective traits. SNPs with significant effects in F2:3 RILs are shown as closed circles. b) Genomic positions of QTL validated in the biparental population. Rectangles represent the 10 chromosomes; horizontal segments indicate the positions along the y-axis (Mb) of polymorphic haplotypes between the parents. Validated QTL are labeled with the associated trait codes and coloured blue or red for positive (trait-increasing) or negative (trait-decreasing) effects, respectively. Non-significant haplotypes are coloured grey, while white regions denote monomorphic intervals. For trait descriptions, refer to **Table 1**

QTL associated with PRL, SRL and LRL mapped to different genomic regions and their effects exhibited distinct temporal dynamics. Effects approached a plateau at 14 dat for PRL and SRL, whereas they increased linearly for LRL (**Fig. S7a**), mirroring the corresponding temporal changes in trait values (**Fig. S7b**).

### Assessing the agronomic relevance of seedling root trait QTL

Using bivariate analyses, we evaluated the effect of each of the lead haplotypes underlying the 109 independent QTL associated with seedling root traits on field-measured agronomic traits, including early plant height (PH_V6), anthesis (MF), final plant height (PH_final), lodging (LO), tillering (TILL), and total dry matter yield (TDMY).

The strongest correlation between haplotype effects on seedling root traits and agronomic traits was observed for PH_V6 (r = 0.48, P < 0.001; **Fig. S8**). Lead haplotypes associated with SRL (n = 21) and LRL (n = 13) explained the largest proportion of genetic variance in PH_V6 (15%), whereas those associated with the remaining seedling root traits accounted for 1% to 12% of the genetic variance in PH_V6 (**Table 3**). Among the 109 lead haplotypes underlying QTL, 14 were significantly associated with PH_V6, and their effects on seedling root traits and PH_V6 were strongly correlated (r = 0.9, P < 0.001, **Fig. 5a**). Of these, four haplotypes were associated with SRL, two each with LRL and TRL, and one each with the remaining seedling root traits (**Fig. 5b**).

**Fig. 5.**
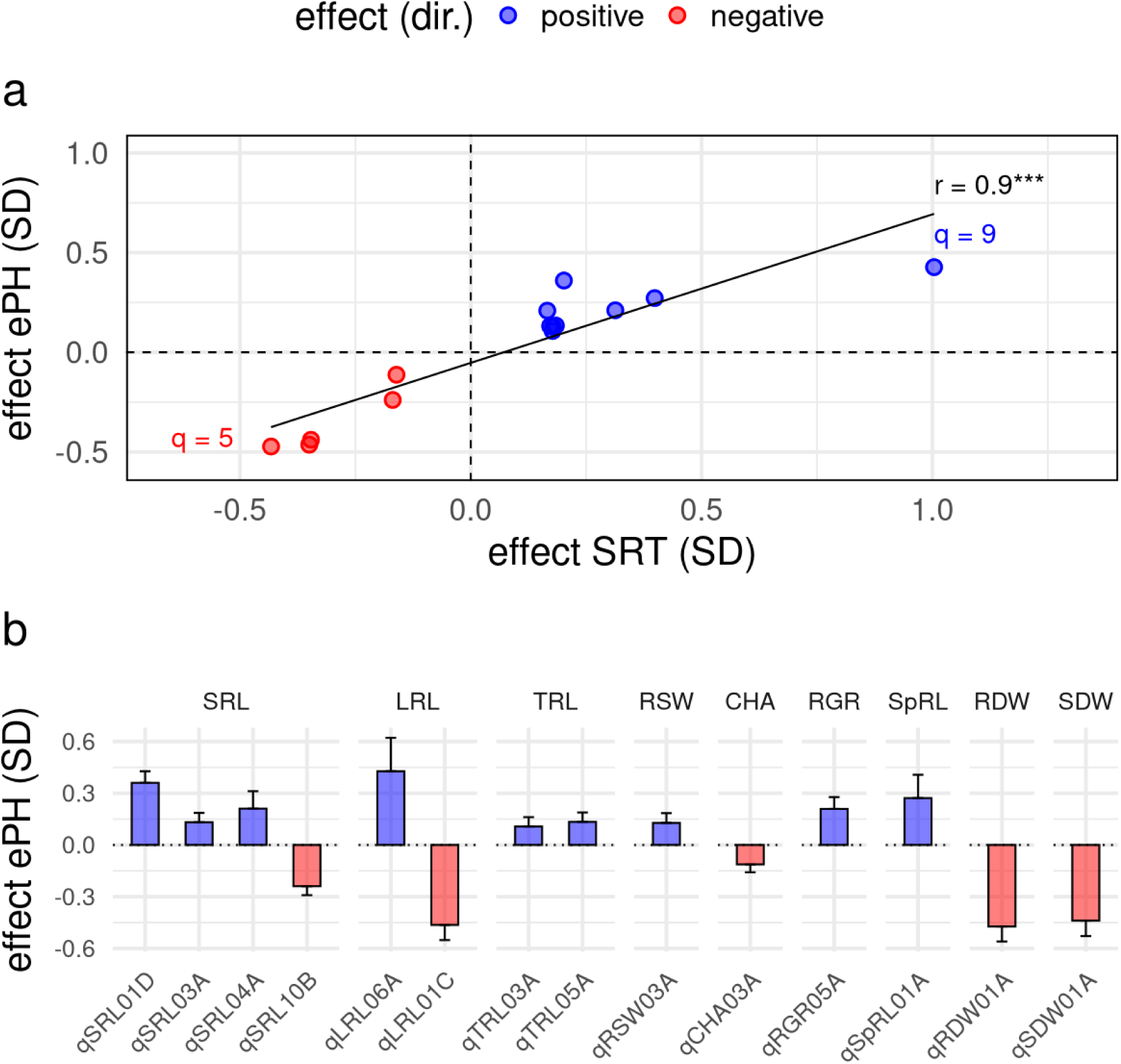
Effect of 14 haplotypes significantly associated with seedling root traits (SRT) and early plant height at stage V6 (ePH). a) Scatterplot of additive haplotype effects on SRT (x-axis) versus ePH (y-axis), expressed in standard deviations (SD) of the respective traits. The solid black line represents a linear regression, with the Pearson correlation coefficient and its significance indicated (*** P < 0.001). Counts of significant haplotypes are indicated in each quadrant. b) Bars represent significant haplotype effects on ePH, expressed as standard deviations (SD) ± standard errors, with panels corresponding to different SRT. In (a) and (b), haplotypes with positive (trait-increasing) and negative (trait-decreasing) effects on SRT and ePH are shown in blue and red, respectively. For trait descriptions, refer to **Table 1**

**Table 3.** Summary information of the effect of lead haplotypes associated with seedling root traits (SRT) on early plant height assessed in the field (PH_V6). Indicated are the Trait code, proportion of genetic variance in PH_V6 explained by the multi-locus model including all lead haplotypes significantly associated with a given SRT [GVE (%)], maximum absolute haplotype effect on PH_V6 expressed in standard deviations [|effect| max (SD)], proportion of haplotypes with concordant, discordant, and significant effects on PH_V6 and the respective SRT [H conc. (%), H disc. (%), H sig. (%)]

| Trait | GVE (%) | \|effect\| max (SD) | H conc. (%) | H disc. (%) | H sig. (%) |
| --- | --- | --- | --- | --- | --- |
| PRL | 1 | 0.10 | 70 | 30 | 0 |
| SRL | 15 | 0.36 | 57 | 43 | 19 |
| LRL | 15 | 0.46 | 77 | 23 | 15 |
| TRL | 11 | 0.46 | 100 | 0 | 60 |
| RSD | 6 | 0.10 | 56 | 44 | 0 |
| RSW | 5 | 0.18 | 85 | 15 | 8 |
| CHA | 3 | 0.29 | 74 | 26 | 11 |
| BAL | 2 | 0.08 | 50 | 50 | 0 |
| RGR | 12 | 0.46 | 64 | 36 | 21 |
| SpRL | 3 | 0.27 | 100 | 0 | 14 |
| RDW | 9 | 0.47 | 43 | 57 | 14 |
| SDW | 11 | 0.44 | 100 | 0 | 14 |

Significant, although weaker, correlations were also detected between haplotype effects on seedling root traits and other agronomic traits except for TDMY (**Fig. S8**). Positive correlations were observed for PH_final and LO, whereas negative correlations were observed for MF, FF, and TILL. 35 of the 109 QTL were also significantly associated with at least one agronomic trait, of which 12 were associated with two or more agronomic traits (**Fig. S9**). Most haplotypes that increased PH_V6 also influenced PH_final significantly in the same direction and were also associated with earlier flowering.

To evaluate the potential of lead haplotypes associated with seedling root traits for improving early plant development in elite germplasm, we assessed their frequency in a panel of 65 flint breeding lines. 18 lead haplotypes associated with increased seedling root trait values, of which seven also significantly affected at least one agronomic trait, were not found in the breeding lines (**Fig. S10a**). Among these, three also increased PH_V6, one promoted earlier flowering, whereas the remaining three exhibited undesirable effects on LO, TILL, or TDMY (**Fig. S10b**).

## Discussion

Optimizing root system architecture is critical for efficient water and nutrient uptake, supporting biomass production while minimizing nutrient losses to deeper soil layers. Therefore, establishing the contribution of seedling root traits to plant performance is essential for developing cold-tolerant maize with enhanced early growth, higher yield potential, and reduced environmental impact in temperate regions (Ojeda-Rivera et al. 2025).

However, our understanding of the genetic regulation and functional role of root traits in maize is limited by the complexity of studying root system architecture (Hochholdinger et al. 2018). High throughput platforms have proven valuable for dissecting the genetic architecture of complex traits by enabling large-scale, non-destructive, and dynamic assessment of plant phenotypes (Xiao et al. 2022). Here, we used the high throughput GrowScreen-PaGe platform to evaluate early root system development under chilling conditions in 815 DH lines derived from two European maize landraces (**Fig. 1, Fig. S1**). Our dataset exhibits high genetic diversity for seedling root and shoot traits and is unprecedented in dimension for maize root studies conducted under chilling conditions, thus providing a powerful and robust foundation for identifying genetic variants influencing seedling root system architecture and evaluating their contribution to field performance.

Due to its tropical origin, maize has limited tolerance to low temperatures, with severe damage occurring below 10°C (Greaves 1996). In the primary root, changes in growth rate, gene expression, and hormone concentrations occur already under mild temperatures around 20°C (Friero et al. 2023), while temperatures of 16°C have been shown to reduce cell elongation and relative elemental growth rate by 40% compared to 25°C (Nagel et al. 2009). The Kemater and Petkuser landraces are adapted to temperate climates and exhibited pronounced genetic variation for early development in both field and controlled environments, with some of the DH lines showing high levels of cold tolerance (Boughazi et al. 2024; Frey et al. 2020; Hoelker et al. 2019). The phenotyping platform used in this study enabled us to apply a temperature regime that can mimic soil temperatures during early maize growth in temperate regions of central Europe, constraining root growth due to chilling while maximizing differentiation among landrace-derived DH lines.

Accordingly, we observed pronounced genetic variation for our target traits with moderate to high heritabilities, which were comparable to or higher than those reported in previous controlled-environment studies of diversity panels (Pace et al. 2015; Sun et al. 2021; Zuffo et al. 2022), breeding lines (Kumar et al. 2012), and exotic introgression lines (Sanchez et al. 2018), highlighting the high quality of the data obtained in the platform and the potential of this landrace-derived plant material as a source of genetic variation for improving root system architecture under chilling conditions (**Table 1**).

We identified several haplotypes significantly associated with these traits, collectively explaining a moderate proportion of the genetic variance, supporting a complex and highly polygenic genetic basis underlying early root trait variation (**Table 2**). These findings are consistent with previous studies conducted in controlled environments (Moussa et al. 2021; Sun et al. 2021; Zuffo et al. 2022), and highlight the importance of large population sizes to achieve sufficient statistical power in GWAS. Most trait-associated haplotypes occurred at intermediate frequencies and exhibited small but consistent effects across landraces. In contrast, haplotypes with moderate to large effects were rare and often specific to a single landrace, potentially reflecting a more recent origin (**Fig. 3, Fig. S6**).

In the biparental population derived from two Kemater DH lines, only common haplotypes could be validated, as none of the rare haplotypes segregated in this population (**Fig. 4**). Because rare variants detected in genome wide analyses are particularly prone to overestimation of effects, validation in an independent population is required to confirm their effect (Göring et al. 2001; Ioannidis et al. 2009). Collectively, the successful identification of both landrace-specific and shared variants underlying early root system architecture variation supports the sampling strategy applied in this study with a large number of DH lines derived from a few pre-selected landraces to characterize the genetic architecture of quantitative traits (Mayer et al. 2017).

QTL associated with seedling root traits have been identified before in maize, but their functional contribution to crop performance has remained poorly understood (Bray and Topp 2018). In this study, a substantial proportion of haplotypes associated with seedling root traits also influenced agronomic traits evaluated across multiple field environments, suggesting pleiotropic or tightly linked effects relevant for plant performance (**Fig. S8**). In line with the importance of the embryonic and early post-embryonic root system described for seedling establishment (Hochholdinger et al. 2004), the strongest correlations among haplotype effects and between traits were observed for early plant height in our study (**Fig. 5, S8**). These results are also consistent with physiological studies indicating that a more developed seedling root system can enhance water and nutrient acquisition and support higher biomass production under chilling conditions (Aroca et al. 2001; Richner et al. 1996). Haplotypes increasing early root and shoot vigour also tended to promote earlier flowering and greater final plant height (**Fig. S9**), in agreement with previous findings (Mayer et al. 2020), most likely resulting from a physiological connection between these traits. However, correlations inferred from marker data should be interpreted with caution, as imperfect linkage disequilibrium (LD) between markers and causal variants can lead to both spurious and reduced detection of genetic correlations among traits (Gianola et al. 2015). Furthermore, pleiotropy can only be distinguished from linkage when a single causal mutation is shown to affect multiple traits, which goes beyond what can be resolved through GWAS (Chebib and Guillaume 2021). While the high marker density used in this study warrants strong LD between haplotypes and causal variants, and the rapid LD decay in this population may favour the detection of pleiotropy over linkage (Schulthess et al. 2017), determining the proportion of truly pleiotropic loci among those identified in the GWAS requires additional evidence beyond the scope of this study.

Determining which root traits should be targeted for adaptation to specific environmental conditions can significantly contribute to the sustainable intensification of agriculture (Lynch 2022). This study focused on the early stages of maize growth under chilling conditions with adequate supply of water and nutrients, during which seedling root system architecture is determined by the primary root, its lateral roots, and seminal roots. Haplotypes associated with both seminal and lateral root length were the most important determinants of early plant height in the field (**Table 3**), consistent with the role of lateral roots in radial uptake of soil resources and of seminal roots in their axial transport (Ahmed et al. 2016). On the other hand, seminal and lateral root length were weakly correlated, did not share overlapping QTL, and exhibited distinct temporal dynamics (**Fig. S4, Fig. S5, Fig.S7**). Lateral root length and relative haplotype effects increased linearly over time, suggesting continued development beyond the observation period. However, most genes influencing embryonic root development do not appear to affect shoot borne root architecture, as exemplified by *rum1*, which regulates early seminal and lateral root initiation but has no effects at later stages (Woll et al. 2005). Additionally, it has been shown that the genetic regulation of lateral root length in shoot borne roots differs between vegetative and reproductive stages (Guffanti et al. 2026). Together, these findings suggest that the loci identified in this study for both seminal and lateral root length are likely to affect only the early stages of root system architecture.

Maintaining the root system represents a significant metabolic cost; therefore, its expansion must be compensated by increased uptake of soil resources, which underlies the context-dependent utility of root system architecture (Lynch et al. 2005). Compared to seminal root length, lateral root length showed stronger phenotypic correlations with early developmental traits and did not exhibit adverse trade-offs with other agronomic traits, supporting early lateral root length as a favourable breeding target. Along the same line, increased lateral root length and density at the seedling stage have been associated with enhanced phosphorus uptake and improved shoot growth under chilling conditions (Hund et al. 2007; Zhu and Lynch 2004), as well as improved seedling survival under drought (Wang et al. 2016; Yu et al. 2024), owing to their relatively low metabolic cost compared with seminal roots.

Recently, it has been demonstrated that domestication and modern breeding indirectly modified root system architecture while improving maize productivity (Ren et al. 2022; Yu et al. 2024). Increasingly challenging climatic conditions call for more targeted selection strategies and the introgression of favourable alleles into elite germplasm, guided by the growing understanding of the genetic regulation and functional role of root traits for crop productivity (Hochholdinger and Yu 2026). Given the highly polygenic basis underlying seedling root trait variation and the complexity of root phenotyping, genomic selection might represent a promising strategy for improving early root system architecture. In this context, high-throughput phenotyping platforms can enable the characterization of large training populations under target environmental conditions, thereby improving prediction accuracy and allowing greater selection intensity.

In addition to enriching our knowledge on the genetic basis of seedling root traits, our study demonstrated that European maize landraces harbor useful genetic variants for improving seedling root system architecture under chilling conditions that are absent from flint breeding lines despite their adaptation to temperate climate conditions (**Fig. S10**). In particular, large effect variants represent promising and accessible targets for marker assisted backcrossing into elite germplasm or as templates for genome editing, given the availability of well-established breeding and molecular tools. Although fine mapping and functional validation of the underlying causal genes are important future steps to confirm their effects prior to introgression, the identified markers, haplotypes, and associated phenotypic effects already provide a solid basis with initial evidence supporting their promising and straightforward implementation in breeding. For haplotypes shared across landraces, concordant effects and validation in a biparental population provided evidence for their stability and potential transferability across genetic backgrounds. Further validation in additional genetic backgrounds and multi environment field trials will assist in confirming their agronomic value (Khaipho-Burch et al. 2023).

## Statements and declarations

### Author Contributions

C.-C.S., M.O., K.N., A.G., and F.G. designed the research and developed ideas, M.O., T.P., D.S, F.G., and C.U. developed and genotyped the plant material, K.N., A.G., C.M, and S.P. performed the phenotyping experiments, F.G. analyzed the phenotypic and genotypic data, CCS and MO acquired funding for the study, F.G., C.-C.S., and K.N. wrote the manuscript, all authors have read and approved the final version of the manuscript; C.-C.S. agrees to serve as the author responsible for contact and to ensure communication.

### Data Availability

All data and materials are available through material transfer agreements upon request.

### Ethical standards

The authors declare that this study complies with the current laws of the countries in which the experiments were performed.

## Acknowledgements

We would like to thank Jonas Lentz, Ann-Katrin Kleinert, Bernd Kastenholz and Sandun Arumayaman for their assistance in the phenotyping experiments.

