## Supplementary Figures for "Genetic analysis of maize seedling root traits under chilling highlights their importance for early field development"

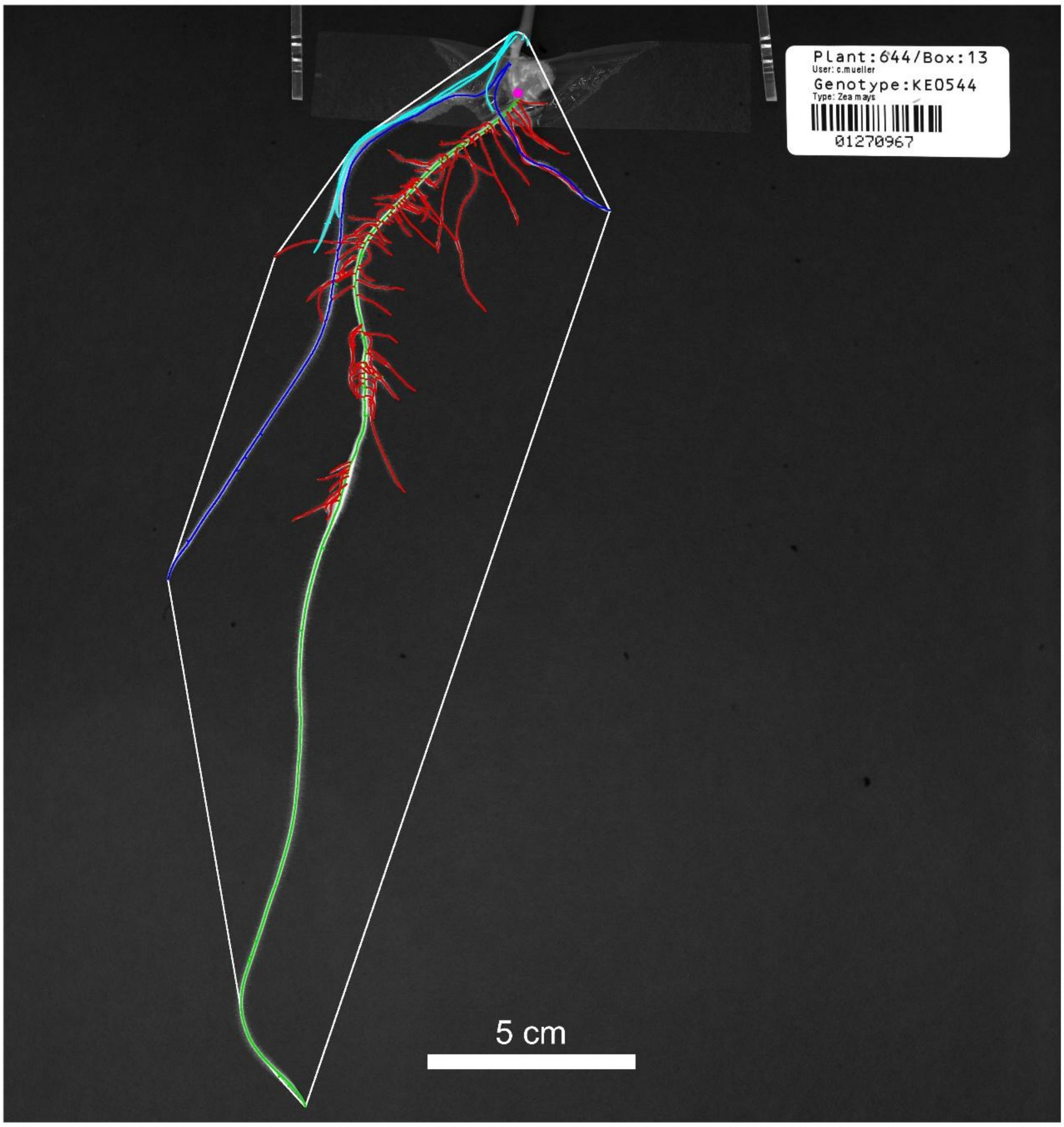


**Fig. S1** Seedling root traits assessment from root pictures.
Analysis of a picture presented in **Fig. 1**, illustrating the seedling root system of a representative plant 14 days after transplantation (dat). Using custom software and manual curation, images were overlaid with a mask distinguishing the different root types: primary (green), seminal (dark blue), crown (turquoise), and lateral roots (red). The white polygon encompassing the root system was used to measure convex hull area, root system width and root system depth. Root traits were extracted automatically from the mask at 5, 7, 9, 12, and 14 dat, while root and shoot dry weights were measured at harvest (14 dat)

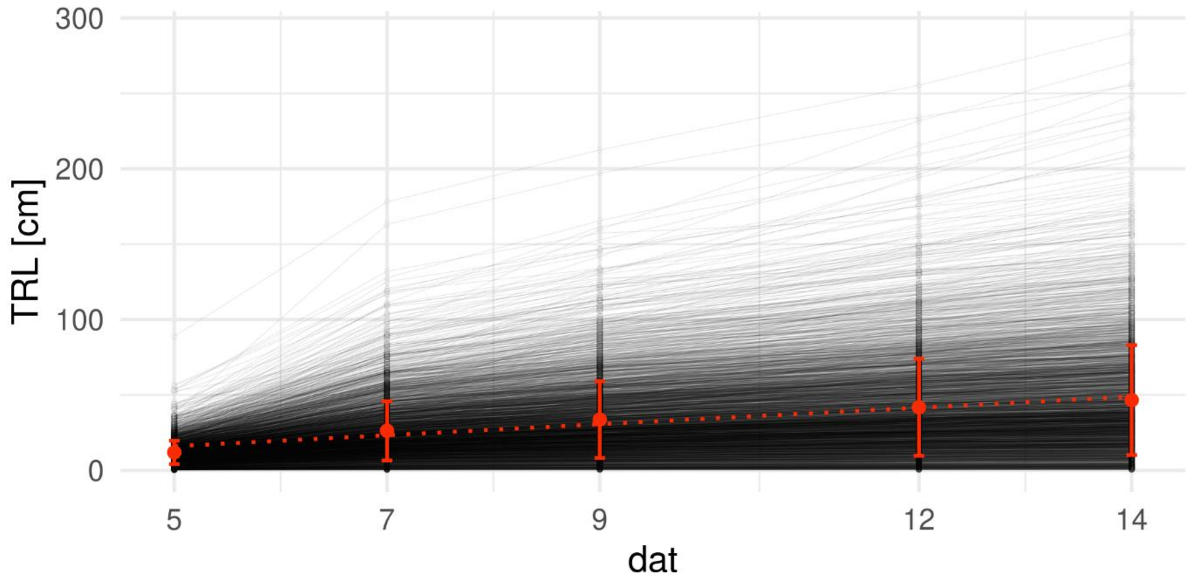


**Fig. S2** Increase in total root length (TRL) over consecutive time points.
TRL of all analyzed plants at 5, 7, 9, 12, and 14 days after transplantation (dat). Black dots connected by lines represent measurements on individual plants. Large red dots indicate mean TRL at each time point, with error bars showing the standard deviation. The dotted line represents a linear regression of TRL over dat

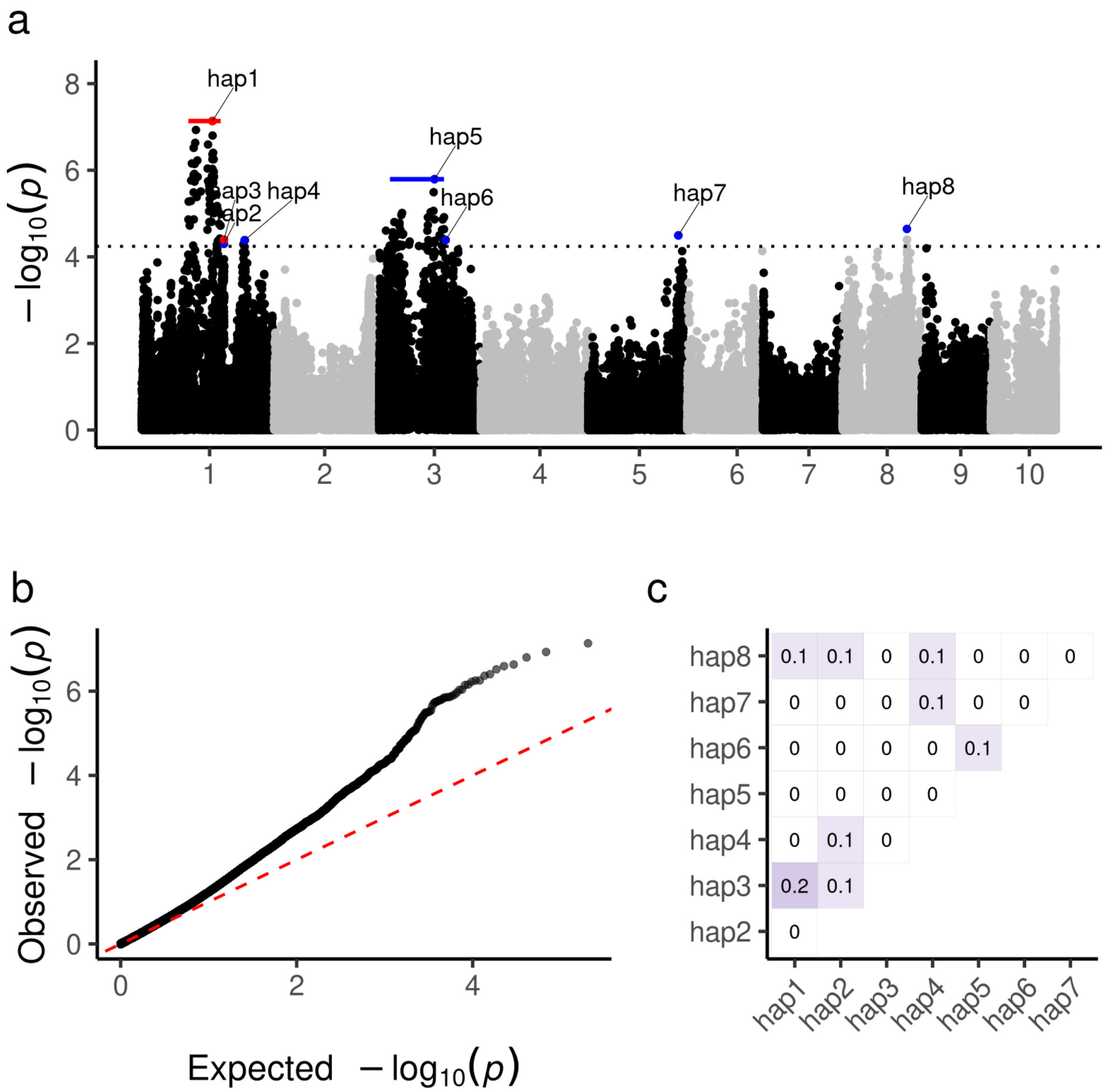


**Fig. S3** Manhattan and quantile-quantile (Q-Q) plots of GWAS results for total root length (TRL).

a) Manhattan plot. Each dot represents a haplotype, plotted by physical position (x-axis) against the significance of its association with TRL ($-\log_{10}(p)$, y-axis). The horizontal dashed line indicates the significance threshold at a 5% false discovery rate. Haplotypes included in the multi-locus model selection are labelled and colored blue or red for positive (trait-increasing) or negative (trait-decreasing) effects on TRL, respectively. Horizontal bars of the same colors indicate the boundaries of genomic regions defined by the first and last significant haplotypes in high linkage disequilibrium (LD) ($r^2 > 0.6$) with these haplotypes.

b) Q-Q plot showing the expected distribution of p-values under the null hypothesis of no association (x-axis) versus the observed distribution (y-axis). The red dashed line indicates the diagonal (y = x).

c) Pairwise LD among the eight haplotypes (hap1-hap8) highlighted in (a), shown as $r^2$ values within each cell and visualized by blue shading. After backward elimination, five of these haplotypes were retained in the multi-locus QTL model, all significantly associated with TRL (hap1, hap4, hap5, hap7, and hap8)

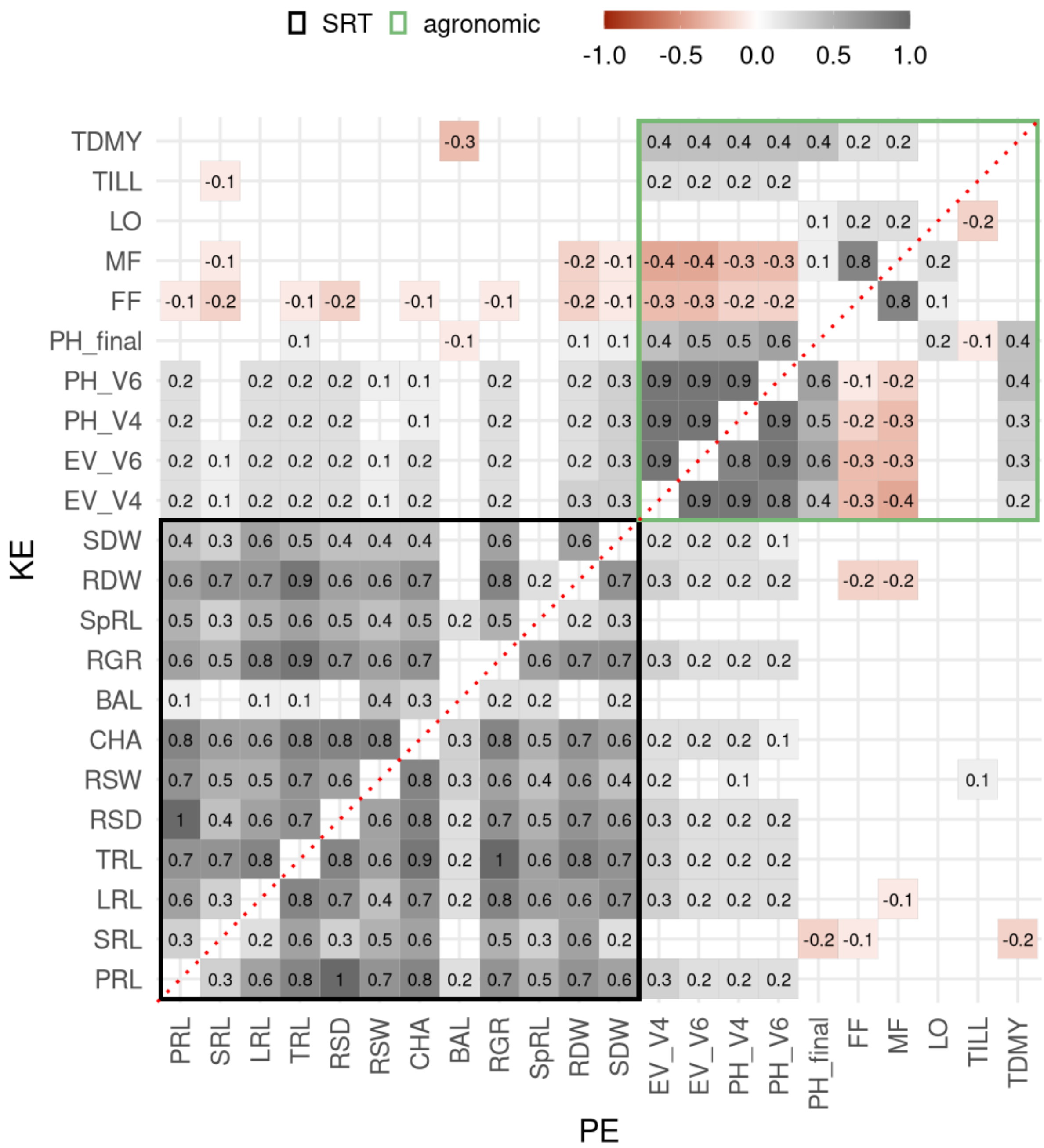


**Fig. S4** Phenotypic correlation of seedling root traits and agronomic traits. Phenotypic Pearson correlation coefficients for twelve seedling root traits (SRT, black rectangle), and ten agronomic traits (green rectangle) within DH lines derived from Kemater (n = 440, above diagonal) and Petkuser (n = 375, below diagonal). The Pearson correlation coefficient is reported within the respective cells and indicated by the colour scale. Only significant correlations (FDR 5%) are displayed.
For trait descriptions, refer to **Table 1**. Agronomic trait codes: EV = early vigour (growth stages V4, V6), PH = plant height (growth stages V4, V6, and final), FF = silking, MF = anthesis, LO = lodging, TILL = tillering, TDMY = total dry matter yield

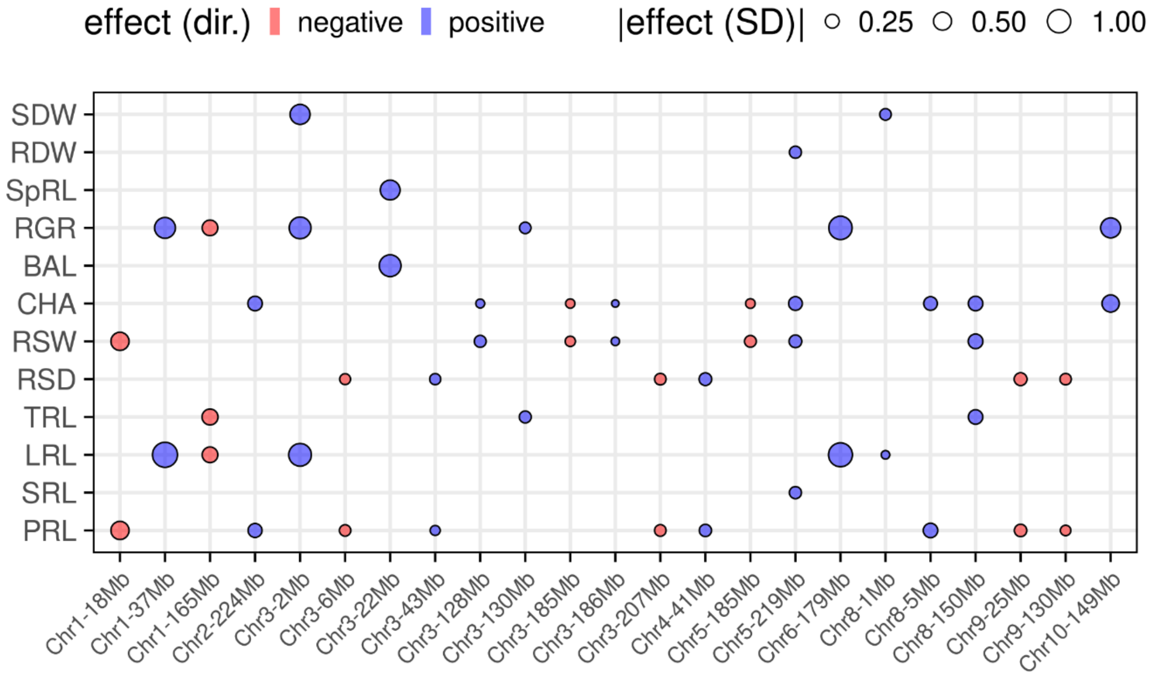


**Fig. S5** QTL associated with multiple seedling root traits (SRT).
Chromosome and position in megabases (x-axis) of QTL associated with multiple SRT. Dots represent the lead haplotypes underlying the QTL, coloured blue or red for positive (trait-increasing) or negative (trait-decreasing) effects, respectively, with size proportional to the additive effect expressed in standard deviations of the respective trait. For trait descriptions, refer to **Table 1**

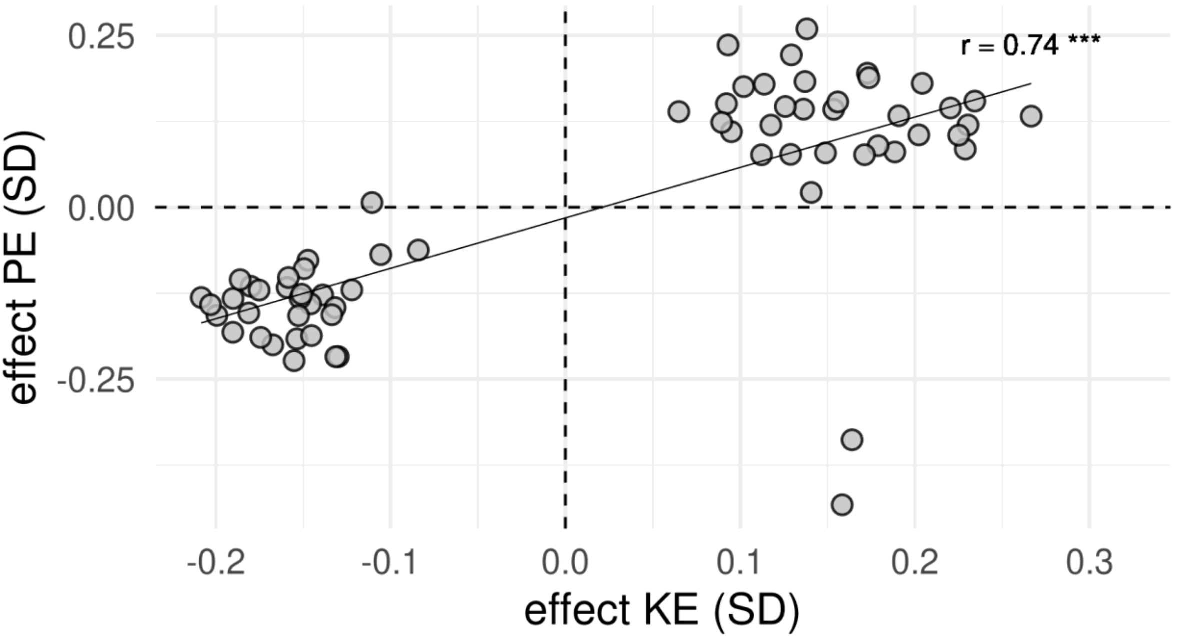


**Fig. S6** Correlations of haplotype effects among landraces.
Scatterplot of the additive haplotype effects estimated within DH lines derived from Kemater (effect KE, x-axis) and Petkuser (effect PE, y-axis), expressed in standard deviations (SD) of the respective traits. The solid black line represents a linear regression, with the Pearson correlation coefficient and its significance indicated (*** $P < 0.001$). Only haplotypes significantly associated with seedling root traits that were present in at least three genotypes within both landraces were included ($n = 66$)

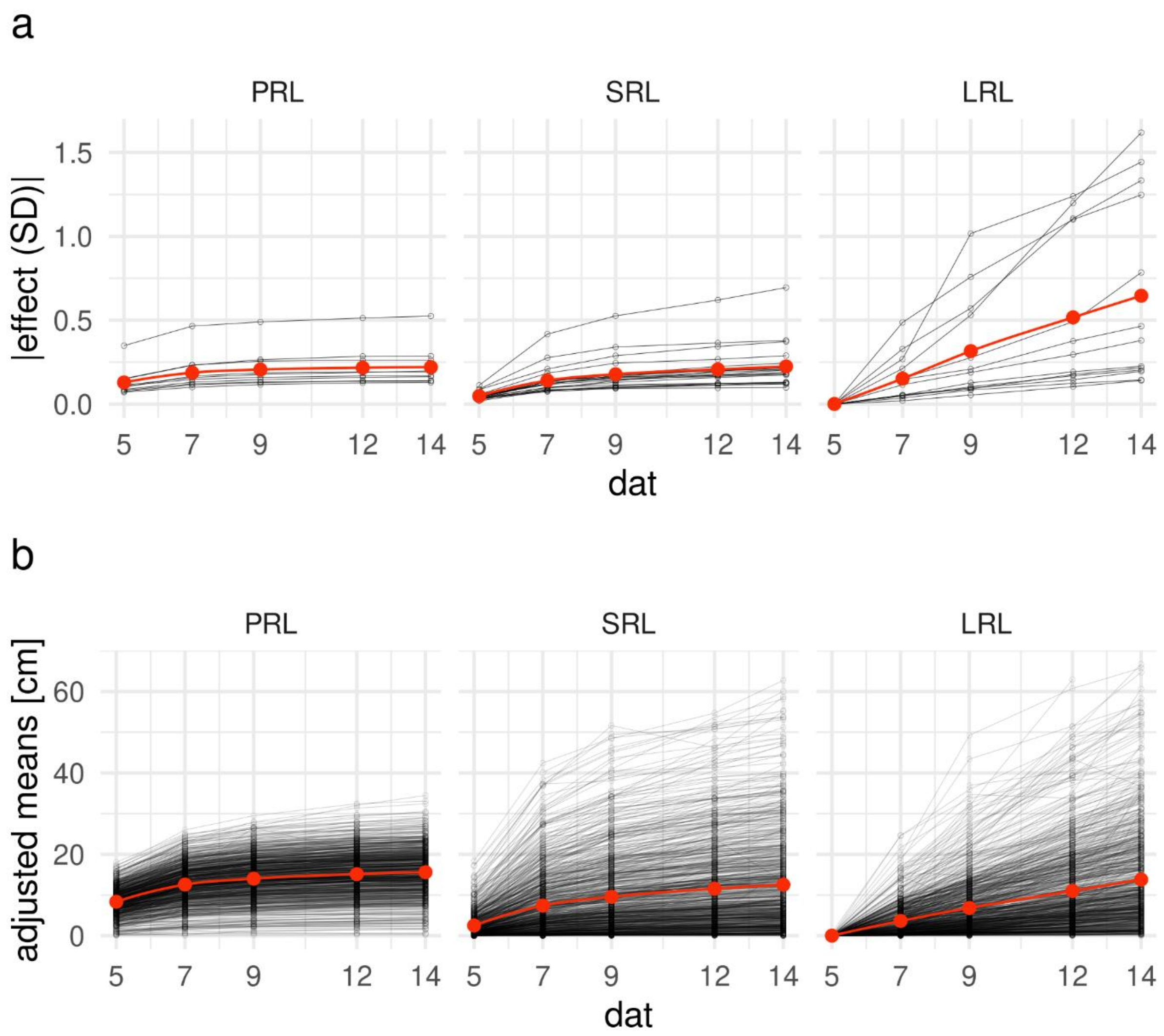


**Fig. S7** Temporal dynamics of haplotype effects and trait variation.
Additive haplotype effects, expressed in standard deviations of the respective trait (a), and adjusted means (b) for primary root length (PRL, left), seminal root length (SRL, center), and lateral root length (LRL, right) at 5, 7, 9, 12, and 14 days after transplantation (dat). Black dots connected by lines represent individual haplotypes in (a) and genotypes in (b). Large red dots indicate mean values at each time point. The solid red line represents a nonlinear fit obtained by local polynomial regression. For trait descriptions, refer to **Table 1**

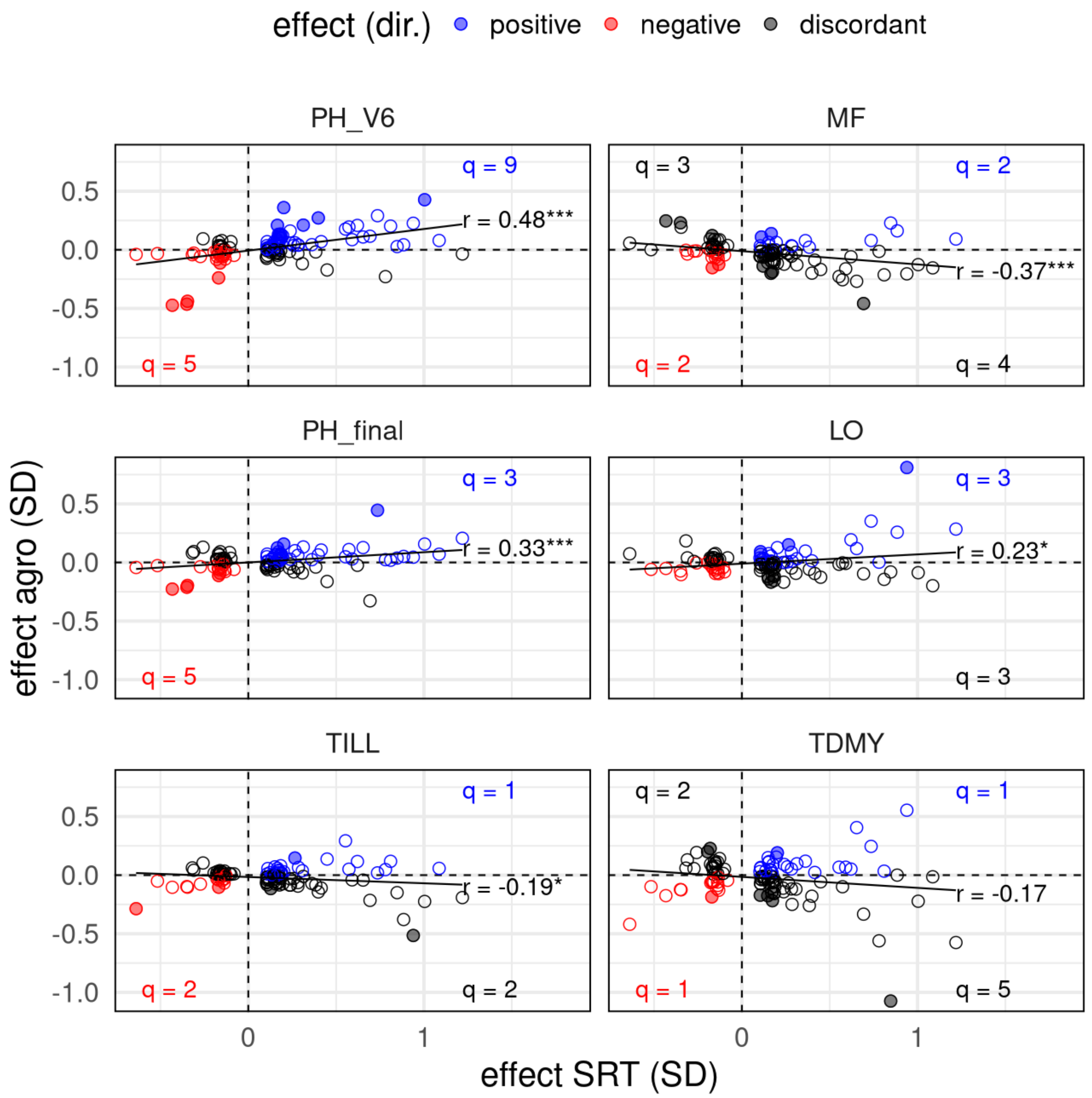


**Fig. S8** Effect of 109 haplotypes associated with seedling root traits (SRT) on agronomic traits.

Scatterplot of additive haplotype effects on SRT (x-axis) versus six agronomic traits (y-axis), expressed in standard deviations of the respective traits. For each agronomic trait (panels), the solid line represents a linear regression, with Pearson correlation coefficients and corresponding significance indicated (*** $P < 0.001$; ** $P < 0.01$; * $P < 0.05$). Haplotypes with positive (trait-increasing), negative (trait-decreasing), and discordant effects on SRT and agronomic traits are shown in blue, red, and black, respectively. Closed and open circles denote haplotypes with significant and non-significant effects on agronomic traits, respectively, with counts of significant haplotypes indicated in each quadrant.

Agronomic trait codes: PH_V6= early plant height (stage V6), MF = anthesis, PH_final = final plant height, LO = lodging, TILL = tillering, TDMY = total dry matter yield

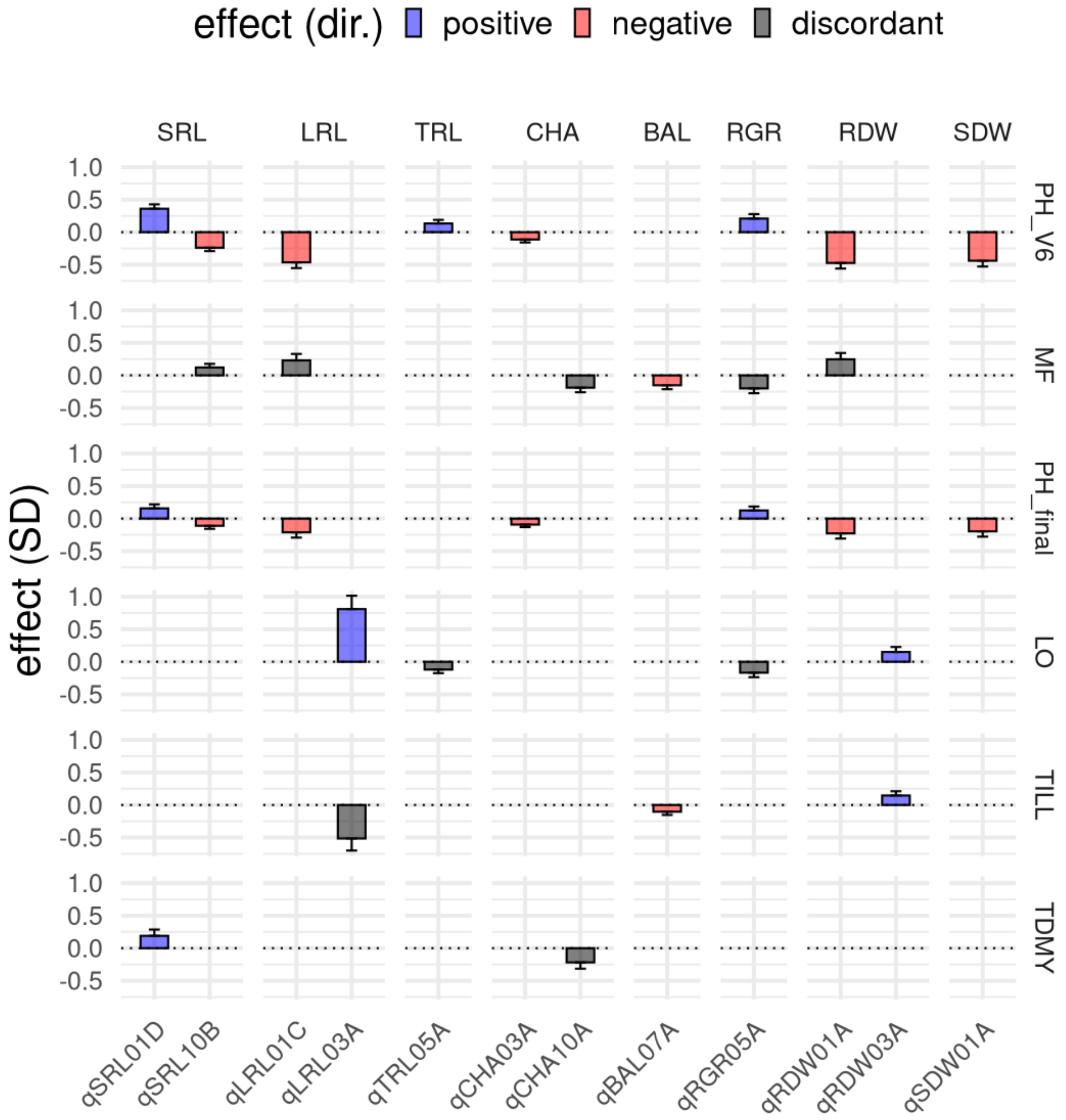


**Fig. S9** Effect of 12 haplotypes associated with seedling root traits (SRT) with significant effects on two or more agronomic traits.

Panels show combinations of SRT (columns) and agronomic traits (rows). Bars represent significant additive haplotype effects expressed in standard deviations (SD) of the respective traits ± standard errors and are colored blue, red, or black if haplotypes exhibited positive (trait-increasing), negative (trait-decreasing), or discordant effects on SRT and agronomic traits, respectively. For trait descriptions, refer to **Table 1.** Agronomic trait codes: PH_V6 = plant height at stage V6, MF = anthesis, PH_final = final plant height, LO = lodging, TILL=tillering, TDMY = total dry matter yield

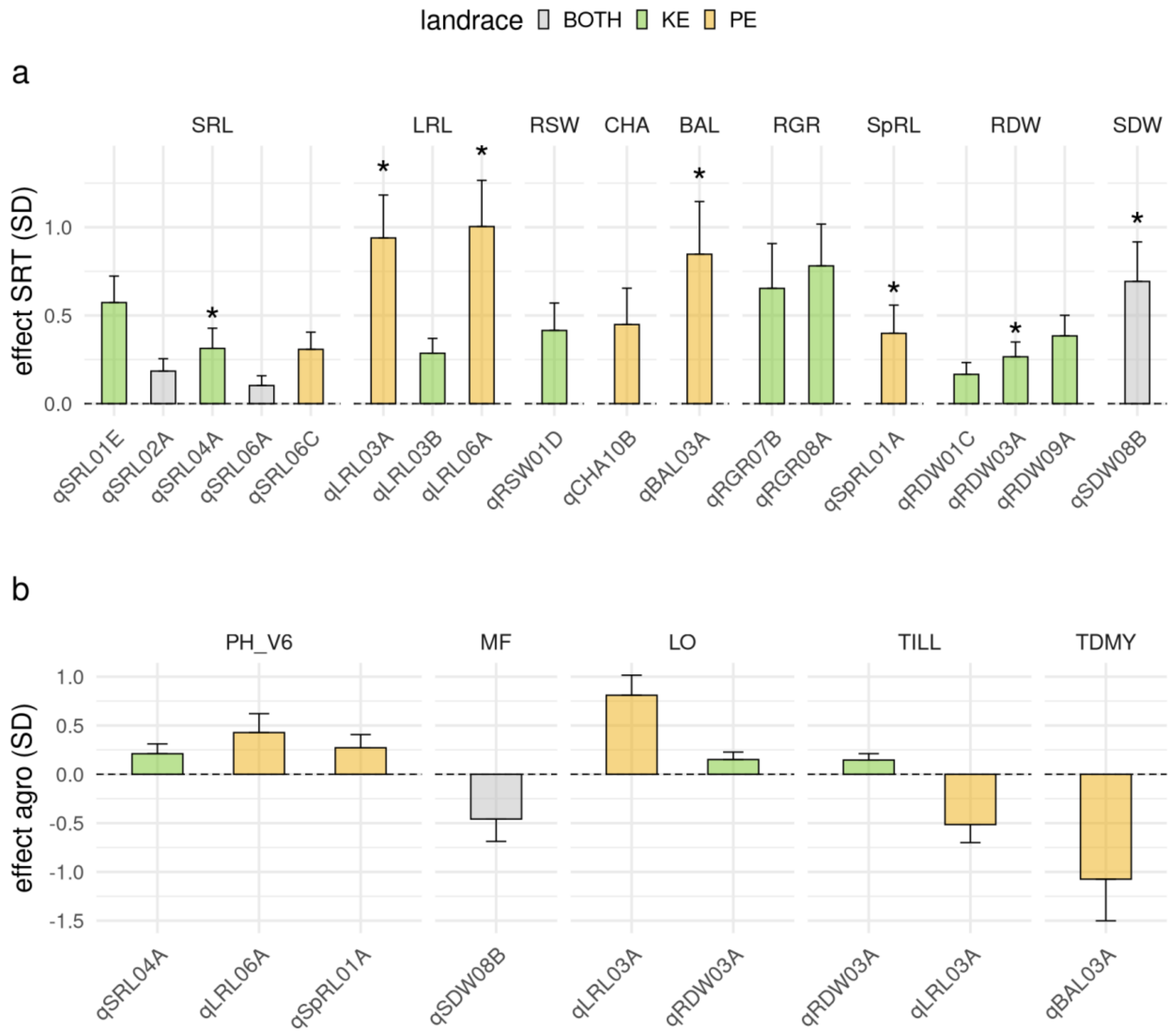


**Fig. S10.** Effects of 18 haplotypes absent from breeding lines with positive effects (trait-increasing) on seedling root traits (SRT).
a) Bars represent significant additive haplotype effects on SRT, expressed in standard deviations (SD) ± standard errors. Haplotypes that also show significant effects on agronomic traits (n = 9) are indicated by asterisks.
b) Bars represent significant additive effects of the haplotypes highlighted in (a) on the respective agronomic traits (panels), expressed in SD ± standard errors.
In (a) and (b), bars are colored grey, green, or yellow to indicate haplotypes present in both landraces, or specific to Kemater (KE) or Petkuser (PE), respectively.
For trait descriptions, refer to **Table 1**. Agronomic trait codes: PH_V6 = plant height at stage V6; MF = anthesis; LO = lodging; TILL = tillering; TDMY = total dry matter yield
